# Engines of change: Transposable element mutation rates are high and vary widely among genotypes and populations of *Daphnia magna*

**DOI:** 10.1101/2020.09.21.307181

**Authors:** Eddie K. H. Ho, E.S. Bellis, Jaclyn Calkins, Jeffrey R. Adrion, Leigh C. Latta, S. Schaack

## Abstract

Transposable elements (TEs) represent a large and dynamic portion of most eukaryotic genomes, yet little is known about their mutation rates or the correspondence between rates and long-term patterns of accrual. We compare TE activity over long and short time periods by quantifying TE profiles and mutation rates (with and without minimizing selection) among 9 genotypes from three populations of *Daphnia magna* sampled along a latitudinal gradient. The patterns of genome-wide variation observed in nature mirror direct estimates of rates and spectra observed in a multi-year laboratory mutation accumulation experiment, where net rates range from -11.98 to 12.79 x 10^-5^ per copy per generation across genotypes. Overall, gains outnumber losses and both types of events are highly deleterious based on comparing lines with and without selection minimized. The rate and spectrum of TE mutations vary widely among genotypes and across TE families/types, even within the same population. We compare TE mutation rates to previously published rates of base substitution, microsatellite mutation, and gene conversion for the same genotypes, and show a correlation only with the latter. Our study provides strong evidence for the notion that TEs represent a highly mutagenic force in the genome. Furthermore, the variation we observe underscores the need to expand the repertoire of mutations studied to include a wider array of mutation types with different underlying mechanisms in order to better understand the evolution of the mutation rate and the ways in which genetic variation is generated genome wide.

## Introduction

It is now known that transposable elements (TEs) make up a significant proportion of the genome in most eukaryotes, and in some cases even represent the majority of the sequence (e.g., Bennetzen and Wang 2014, Canapa et al. 2015, Sotero-Caio et al. 2016, Bourque et al. 2018). Although commonly referred to as ‘junk’ or genomic ‘parasites’, and therefore masked in genomic analyses in favor of focusing on genic regions, the importance of TEs is gaining wider appreciation and the repetitive landscape of the genome is no longer completely ignored (Goerner-Potvin and Bourque 2018, Lanciano and Cristofari 2020). Notably, there are now many examples of TEs performing functional roles in the host genome (e.g., van’t Hof et al. 2016) and recent work has cited their role in biological processes such as adaptation and speciation (Schrader and Schmitz 2018, Serrato-Capuchina and Matute 2018). The influence of TEs at the genomic level, whether direct (e.g., contributing new coding or regulatory sequences; Joly-Lopez and Bureau 2018) or indirect (e.g., changing the epigenomic landscape in the host genome; Pehrsson et al. 2019, Choi and Lee 2020), can be highly influential.

Because TEs are mobile and far outnumber ‘regular’ protein-coding genes in most eukaryotic genomes, elucidating their patterns of replication, transposition, and excision/deletion is a major task that spans subdisciplines from molecular biology to population genetics (Hickman and Dyda 2015, Bourgeois and Boissinot 2019). The dynamics of TE proliferation includes 1) how TEs jump between lineages initially (horizontal transfer of TEs; HTT; Zhang et al. 2020), 2) differential success among TE families in various host lineages (e.g., Lu et al. 2017), and 3) when and where TEs are repressed and/or go extinct, and become resurrected (e.g., Blumenstiel 2019). Indeed, the idea that genomes are like habitats and that TEs are like individuals (and TE families like species) has gained popularity as a way of better characterizing the complexities of TE movement in different host genomes (e.g., Kremer et al. 2020) and the notion that TEs and their host genomes co-evolve is now widely acknowledged (Koonin et al. 2020). Rather than thinking of TEs as simply selfish or parasitic DNA, the emerging body of work on TEs reveals that their mutagenic potential can have a range of outcomes from beneficial to neutral to deleterious (Song and Schaack 2018).

Ultimately, the net TE activity observed in a lineage is the product of the intrinsic mutational properties of the TEs, combined with the host genome’s cellular and genomic defense system, which is then governed (over evolutionary time scales) by population genetic factors-- i.e., the relative strength of selection and genetic drift. An important question is to what degree the baseline mutational inputs are altered in natural populations, once other evolutionary forces act and interact. If TEs, like other categories of mutation, are on average slightly deleterious, they should be purged in lineages where selection can operate efficiently and never accumulate to high copy number. On the other hand, if effective population sizes or recombination rates are low, selection may not act efficiently, and TEs could accumulate. Comparing TE dynamics in the lab versus in nature is critical for unveiling the individual influence of mutation, selection, and drift on this portion of the genome (e.g., Adrion et al. 2017). Furthermore, comparing TE dynamics among closely-related lineages reveals how the mutational process and/or evolutionary constraints vary within and between genotypes, populations, and species. Finally, different types of mutations (e.g., TEs versus base substitutions) may differ in the ways they are shaped by evolutionary forces, making an exploration of this contrast among mutation categories important.

Here, we compare patterns of TE activity over short time periods (using an mutation accumulation [MA] experiment, where selection is minimized) to patterns of long-term accumulation (by comparing genotypes across populations and between congeners) using *Daphnia*. *Daphnia* are an excellent model organism for studying TEs (and mutations, more broadly) because they can reproduce asexually, removing the complicating influence of meiosis and sex on proliferation. *Daphnia* are aquatic microcrustaceans (Order: Cladocera) long used in ecological and toxicological studies, but which have more recently become the focus of evolutionary and genomic research as well (Schaack 2008). While *D. magna* and its congener, *D. pulex,* appear extremely similar in morphology, physiology, behavior, and life-history, they differ in genome size (Vergilino et al. 2009, Jalal et al. 2013) and estimates of effective population size (N_e_; Haag et al. 2009). In this study, we quantify the TE profiles of 9 starting genotypes sampled from three populations of *D. magna* across a latitudinal gradient (Finland, German, and Israel). We use those same genotypes to perform a multi-year MA experiment to directly estimate rates of gains and losses for all known TEs. We also compare our results to *D. pulex*, for which some similar data are available, and to mutation rates for other types of mutation (i.e., microsatellite and base substitution rates) that have been measured in *D. magna* previously (Ho et al. 2019, 2020).

If the patterns observed over short and long time periods are similar (e.g., the most active TEs currently are the most abundant in the genome), it suggests that mutation plays the major role in determining TE content and other evolutionary forces are weak. In contrast, if the patterns of accrued TEs over long time periods do not mirror the observations in real time, it suggests that evolutionary forces other than mutation (natural selection and genetic drift) shape TE content significantly. Patterns of long-term TE accumulation can be measured in several ways: abundance and diversity of TEs present in the genome, insertion site polymorphism (ISP) among lineages, and mean pairwise divergences (MPDs) of copies of each TE family, where lower values are assumed to represent more recent activity, as copies will not have diverged yet due to the accumulation of point mutations. Direct observations of TE movement in real-time using MA experiments represent the gold standard for accurate rate estimates (reviewed in Katju and Bergthorsson 2019), and have only been performed in a few species for mutations involving mobile DNA (McIlroy et al. *in prep*). In these experiments, descendent lineages are propagated via single-progeny descent from a known ancestor to minimize natural selection, lines are sequences and compared to count the number of events per copy per generation and calculate rates. Importantly, while there are two kinds of events that can be scored for a particular TE copy-- gains and losses-- there are a number of ways by which these two events can occur even in asexually-reproducing lineages. New TE copies can result from insertions (transposition or retrotransposition), duplication events, polyploidization, DNA repair, gene conversion events, and/or ectopic recombination. Similarly, loss of a TE can be due to excision (although not all elements are capable of excision [e.g., Class 1 retroelements]), deletions (if a TE was present in a deleted region), gene conversion, or ectopic recombination events. In the vast majority of cases, the exact mechanism of gain or loss is not known. The likelihood of gain and loss via these different mechanisms may vary, for example among sites that are initially unoccupied, heterozygous, or homozygous for a TE.

In addition to contrasting the real-time rates and spectrum of TE mutations with the long-term patterns of TE accrual in the genome, our investigation into the levels of intra- and interspecific variation in TE dynamics reveals the degree to which population genetic factors influence such patterns in different environments. Specifically, low levels of intra- and interspecific variation (e.g., few differences between congeners) would be predicted if effective population size as it is often conceived of *at the species level* governs TE dynamics (Lynch and Conery 2003). Alternatively, if local environments, interdemic variation, or annual bottlenecks result in major differences among populations in TE dynamics or accumulation, it would support the notion that population genetic parameters are not as important at the species level, but instead act more locally. Finally, our comparison of TE mutation rates to the estimates from other types of mutation provides insight into the relative importance of DNA damage versus DNA repair mechanisms in determining the mutation rate genome-wide. That is, if TE mutation rates across genotypes are highly correlated with base substitution mutation rates, it would indicate that the DNA repair plays a major role in determining rates of mutation. In contrast, if rates depend more on the unique mechanisms that produce new TE insertions versus, for example, base substitutions, rates would not be expected to correlate positively. Investigations of correlations among mutation rates across mutation types are sorely lacking (but see Ananda et al. 2011), and a lack of relationship could have major implications for the generalization of mutation rate estimates in biological models.

## Results

We investigated genome-wide patterns of TE accrual and mobilization across two scales: time (long- and short-term) and space (among lineages within and between species). To assess the long-term patterns of accrual of TEs we surveyed the whole genome of 9 genotypes of *D. magna* from three populations and characterized the TE content using three metrics: 1) overall abundance and diversity, 2) insertion site polymorphism, and 3) mean pairwise divergence among copies in each family or superfamily. To quantify real-time patterns of mobility, we directly estimate TE mutation rates (gains and losses; Figure 1) based on events observed during a multi-year MA experiment initiated from each of these genotypes, where descendant lines were either propagated via single-progeny descent (to minimize selection) or maintained at large population sizes. We examine intra- and interspecific variation by comparing our results from *D. magna* along the latitudinal gradient from which the starting genotypes were collected, and by comparing with the congener, *D. pulex*, when possible. Lastly, we compare TE mutation rates from *D. magna* to mutation and gene conversion rates estimated using the same genotypes to see if patterns of TE rate variation covary with other mutational processes that generate or remove genetic changes.

**Figure 1.**
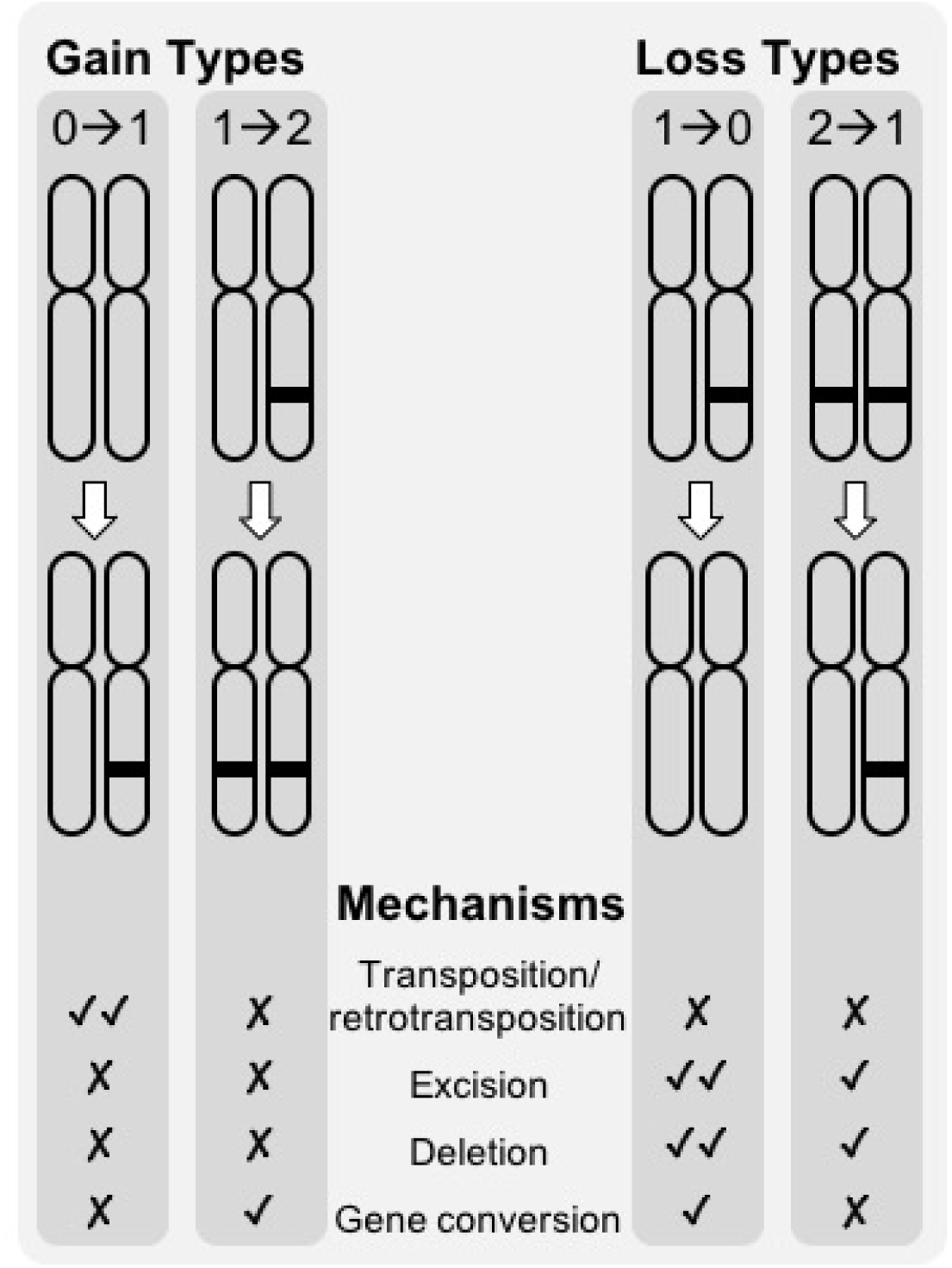
Different mechanisms of loss and gain most likely explain the four categories of loss and gain of TEs at a given locus (0-->1, 1-->2, 1-->0, and 2-->1) that are can occur in asexually-reproducing organisms like *Daphnia magna*. For each type of gain or loss, check marks indicate the relative likelihood (or relative likelihood) of a given mechanism and X marks indicate a particular mechanism cannot produce that type of gain or loss.

### Long-term patterns of TE accrual in Daphnia

To assess variation in patterns of TE accumulation over long time periods, we quantified the genome-wide TE content across 9 genotypes from three populations of *D. magna* along a latitudinal gradient (Figure 2A) and compared it to the congener, *D. pulex*. Among genotypes and populations of *D. magna,* the abundance and diversity of TEs in the genome is very similar, although there are fewer TEs overall than in *D. pulex* (Figure 2B; Table 1). This is true regardless of approach (read mapping versus repeat masking; see Methods and Supplemental Results; Table S3), but it is important to note that estimates of abundance differ by more than a factor of two depending on the method.

**Figure 2.**
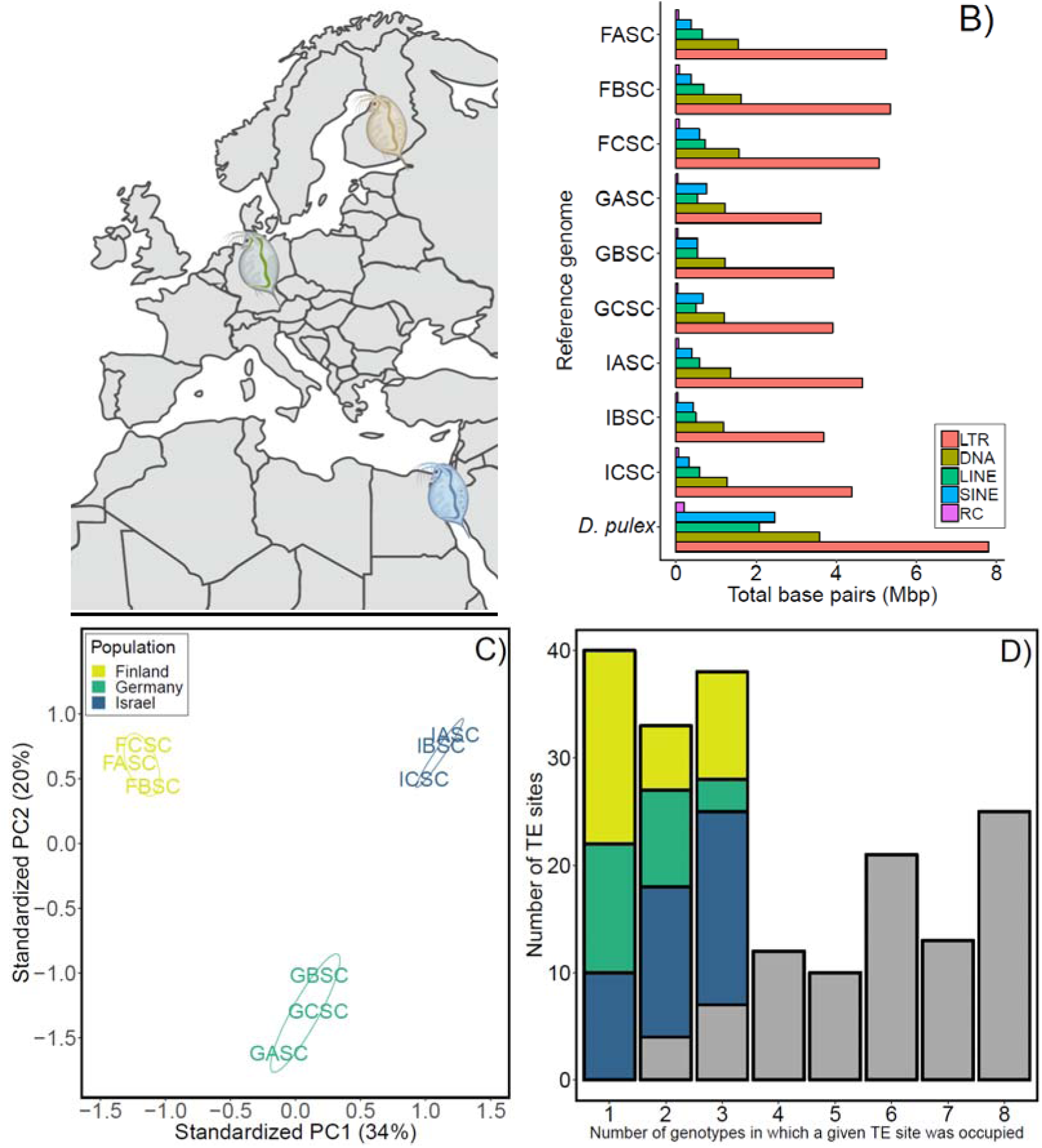
Transposable element profiles of *Daphnia magna* for nine genotypes collected from (A) three populations (Finland [FASC, FBSC, FCSC], Germany [GASC, GBSC, GCSC], and Israel [IASC, IBSC, ICSC). (B) Abundance and diversity (in millions of bp [Mbp] per type of TE (<u>L</u>ong <u>T</u>erminal <u>R</u>epeats [LTR], <u>DNA</u> transposons [DNA], <u>L</u>ong <u>I</u>nterspersed <u>N</u>uclear <u>E</u>lements [LINE], <u>S</u>hort <u>I</u>nterspersed <u>N</u>uclear <u>E</u>lements [SINE], and <u>R</u>olling <u>C</u>ircle elements [RC]) compared to *D. pulex* (reference genome; PA42 [BioProject: PRJEB14656]). (C) Principal Component Analysis shows populations differ based on the presence/absence of TEs across 192 polymorphic sites. (D) Number of polymorphic TE sites occupied; the left bar (x = 1) is the number of singletons (sites occupied in only one genotype), colored portions of bars in x = 2 and x = 3 represent sites occupied in 2 and 3 genotypes, respectively, when from the same population. Grey portions of each bar represent the number of sites that were occupied in ≥2 genotypes that were not population-specific. FASC was used as the reference assembly for (C) and (D); see Supplemental Results.

**Table 1.**
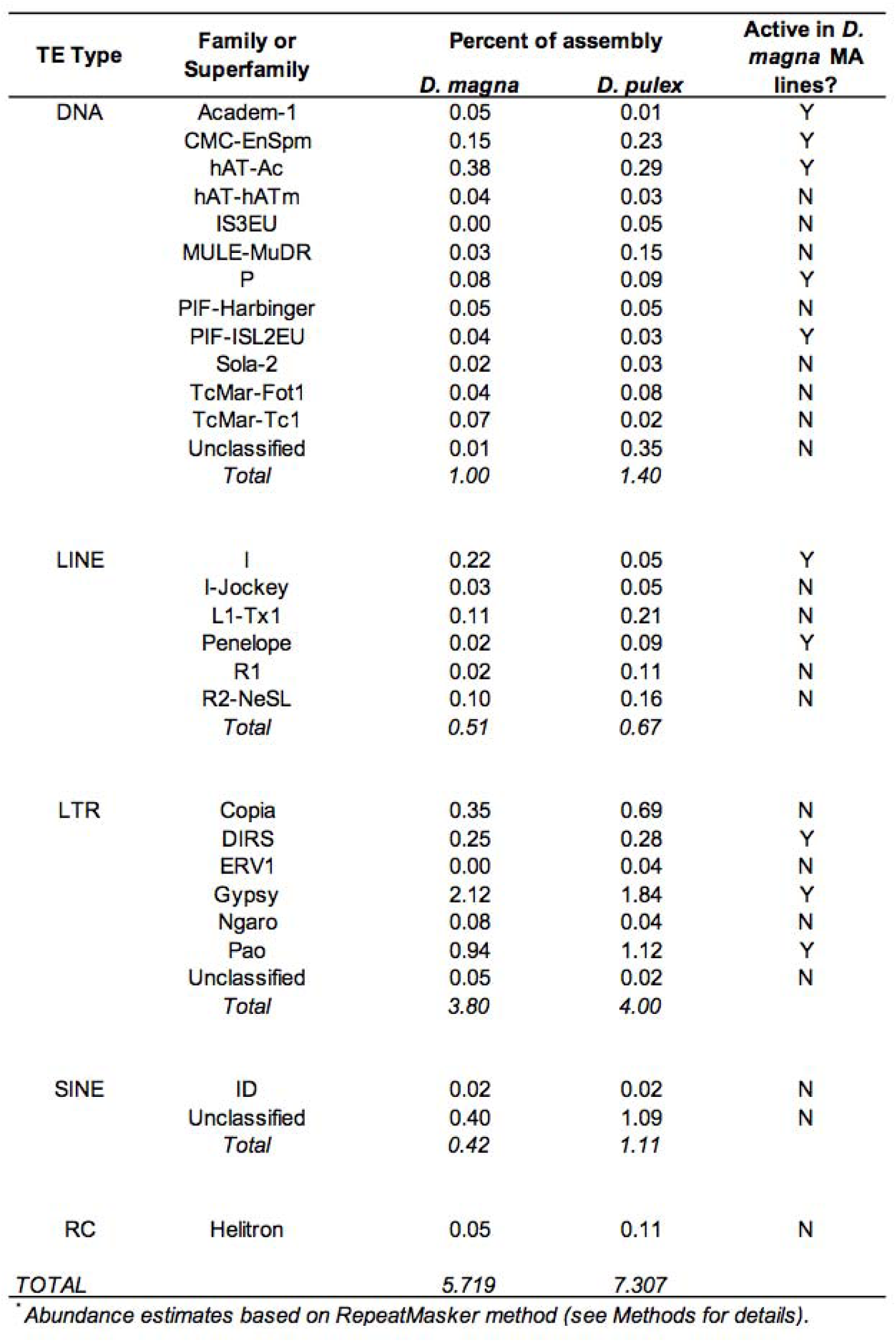
Abundance of TE types by family or superfamily shared by *D. magna* (averaged across nine genotypes) and *D. pulex* (PA42 [PRJNA307976]).

| TE Type | Family or Superfamily | Percent of assembly |  | Active in *D. magna* MA lines? |
| --- | --- | --- | --- | --- |
|  |  | *D. magna* | *D. pulex* |  |
| DNA | Academ-1 | 0.05 | 0.01 | Y |
|  | CMC-EnSpm | 0.15 | 0.23 | Y |
|  | hAT-Ac | 0.38 | 0.29 | Y |
|  | hAT-hATm | 0.04 | 0.03 | N |
|  | IS3EU | 0.00 | 0.05 | N |
|  | MULE-MuDR | 0.03 | 0.15 | N |
|  | P | 0.08 | 0.09 | Y |
|  | PIF-Harbinger | 0.05 | 0.05 | N |
|  | PIF-ISL2EU | 0.04 | 0.03 | Y |
|  | Sola-2 | 0.02 | 0.03 | N |
|  | TcMar-Fot1 | 0.04 | 0.08 | N |
|  | TcMar-Tc1 | 0.07 | 0.02 | N |
|  | Unclassified | 0.01 | 0.35 | N |
|  | *Total* | *1.00* | *1.40* |  |
| LINE | I | 0.22 | 0.05 | Y |
|  | I-Jockey | 0.03 | 0.05 | N |
|  | L1-Tx1 | 0.11 | 0.21 | N |
|  | Penelope | 0.02 | 0.09 | Y |
|  | R1 | 0.02 | 0.11 | N |
|  | R2-NeSL | 0.10 | 0.16 | N |
|  | *Total* | *0.51* | *0.67* |  |
| LTR | Copia | 0.35 | 0.69 | N |
|  | DIRS | 0.25 | 0.28 | Y |
|  | ERV1 | 0.00 | 0.04 | N |
|  | Gypsy | 2.12 | 1.84 | Y |
|  | Ngaro | 0.08 | 0.04 | N |
|  | Pao | 0.94 | 1.12 | Y |
|  | Unclassified | 0.05 | 0.02 | N |
|  | *Total* | *3.80* | *4.00* |  |
| SINE | ID | 0.02 | 0.02 | N |
|  | Unclassified | 0.40 | 1.09 | N |
|  | *Total* | *0.42* | *1.11* |  |
| RC | Helitron | 0.05 | 0.11 | N |
| **TOTAL** |  | **5.719** | **7.307** |  |

\* Abundance estimates based on RepeatMasker method (see Methods for details).

Using read-mapping, 16% of the *D. magna* genome is TEs, whereas repeat masking (which is more common practice) estimates 6% of the genome is TEs (the latter method is used for estimates in Figure 2B and where specified; but see Table S4 for estimates using both methods). Although we recommend using a read-mapping approach for accuracy, as it likely is less prone to bias introduced based on assembly quality, repeat masking is the more commonly used method (see Supplemental Results) and provides the opportunity for inter- as well as intraspecific comparisons here. In *Daphnia*, LTR retrotransposons are the most common type of TEs, with the Gypsy superfamily being the most abundant, regardless of species (Table 1) or method (Table S4). All other categories of TEs (DNA transposons, LINEs, SINEs, and RCs), constitute less than 2% of the genome, although DNA transposons are still highly diverse with 18 different families identified (Table S4). While the relative abundance of most major TE categories is similar between *D. magna* and *pulex*, 7 and 19 families of DNA transposons identified in the genome are specific to *D. magna* and *D. pulex*, respectively. In sum, major differences in abundance and diversity are not readily apparent among genotypes and populations within *D. magna*, but exist between congeners (Figure 2B).

Despite the similarities in abundance and diversity observed within *D. magna*, the TE profiles of genotypes from each of the three populations (Finland, Germany, and Israel) can be easily distinguished based on patterns of insertion site polymorphism using PCA (Figure 2C and Figure S1; presence/absence scored for 192 out of 1442 polymorphic sites). The first and second principal components explained 34% and 20% of the variance, respectively. Whether a particular site is occupied by a TE is determined by events either at the chromosome level (e.g., insertions/deletions, gains/losses due to gene conversion, or differential rates of recombination) or at the individual/population level (e.g., frequency of sexual reproduction in cyclical parthenogens or the strength of selection against new insertions). Depending on the assembly used as the reference (see Supplemental Results), we identified up to 1903 TE sites, of which up to 16% were polymorphic across the nine genotypes (Table S5). On average, we find 19% of TE insertion polymorphisms (TIPs) are specific to a single genotype (i.e., singletons) and an additional 29% of TIPs are specific to a single population (color bars in Figure 2D). Across all TE families, there are typically more singletons found in genotypes from Finland (Figure S2). In contrast, Germany not only exhibited the fewest singletons, but also the fewest population-specific TIPs across all TE types (Table S6; Figure S3).

Estimates of mean pairwise divergence (MPD) among copies for each TE family/superfamily are often used as a proxy for determining when TEs may have invaded and initially been active in the genome; lower values are inferred to mean more recent invasion/activity, because there has been less time to accrue point mutations among copies. While there was a wide range of MPDs observed across superfamilies (14.6% to 30.8%; Table S7A), these cannot be used to accurately estimate the time since invasion/activity for a particular type of TE unless subfamilies, families, and superfamilies are reliably distinguishable. Instead, we looked between species and across genotypes to see if the average MPDs varied (either for TEs known to be active currently [see next section] or for those where no activity has been observed) and there is no difference (Table S7B and C; Figures S4A-L), nor is there a relationship between MPDs across genotypes based on their previously reported base substitution mutation rates (Ho et al. 2020; Figure S5).

In sum, the long-term patterns of TE accumulation within *D. magna* do not exhibit much intraspecific variation in terms of TE abundance or divergence, but do yield a distinctive profile for each population. Overall, these patterns result in a lower genome-wide TE load for *D. magna* than that found in the cosmopolitan congener, *D. pulex*, despite the former having a larger genome (as measured by flow cytometry, *D. magna* = 0.30 pg and *D. pulex* C value = 0.23 pg [Jalal et al. 2016]).

### Real-time rates of TE loss and gain with and without selection

We used mutation accumulation (MA) experiments initiated from each of the 9 genotypes of *D. magna* from each of the three populations to estimate overall (Table 2) and family-specific TE mutation rates (Table 3). Rates of gain across MA lines ranged from 0 to 22.6 x 10^-5^ per copy per generation with a mean rate of 1.39 x 10^-5^ /copy/gen (95% CI: 0.41 x 10^-5^– 2.66 x 10^-5^) and loss rates ranged from 0 to 31.8 x 10^-5^ /copy/gen with a mean of 1.70 x 10^-5^ /copy/gen (95% CI: 0.53 x 10^-5^– 3.23 x 10^-5^; Table S8).

**Table 2.** Number of events and mean TE mutation rates (per copy per generation, including 95% confidence intervals [CI] for gains and losses based on whole genome sequence data from 62 *Daphnia magna* mutation accumulation [MA] and extant controls lines [EC; large population] descended from 9 starting genotypes collected from Finland, Germany, and Israel.

|  | Mutation Accumulation Lines |  |  |  | Extant Control Lines |  |  |  |
| --- | --- | --- | --- | --- | --- | --- | --- | --- |
| | Mutation rate ($10^{-5}$) per copy/ per generation | | | | Mutation rate ($10^{-5}$) per copy/ per generation | | | |
|  | Number of events | Mean | Lower CI | Upper CI | Number of events | Mean | Lower CI | Upper CI |
| Gain (all kinds) | 67 | 1.39 | 0.41 | 2.66 | 2 | 0.009 | 0 | 0.024 |
| 0-->1 gain | 62 | 1.17 | 0.23 | 2.42 | 1 | 0.003 | 0 | 0.008 |
| 1-->2 gain | 5 | 0.22 | 0 | 0.63 | 1 | 0.007 | 0 | 0.019 |
| Loss (all kinds) | 28 | 1.7 | 0.53 | 3.23 | 9 | 0.70 | 0.020 | 1.87 |
| 2-->1 loss | 2 | 0.04 | 0 | 0.09 | 1 | 0.11 | 0 | 0.34 |
| 1-->0 loss | 26 | 1.67 | 0.46 | 3.21 | 8 | 0.58 | 0.020 | 1.65 |
| Total | 95 | 3.09 | 1.37 | 5.14 | 11 | 0.71 | 0.027 | 1.88 |

\*Confidence intervals estimated using by bootstrapping across MA lines 10000 times.

**Table 3.** Number of events and rates (plus 95% confidence intervals [CI]) of gain and loss (per copy per generation) for each TE superfamily in which events were observed averaged across all MA lines. Gains and losses based on whole genome sequence data from *Daphnia magna* mutation accumulation lines descended from 9 starting genotypes collected from Finland, Germany, and Israel.

| Type | Family/<br>Superfamily | Gains |  |  |  | Losses |  |  |  |
| --- | --- | --- | --- | --- | --- | --- | --- | --- | --- |
| | | Number of events | Mean Rate ($10^{-5}$) | Lower CI | Upper CI | Number of events | Mean Rate ($10^{-5}$) | Lower CI | Upper CI |
| DNA | *Academ-1* | 1 | 4.9 | 0.0 | 14.6 | 0 | - | - | - |
|  | *CMC-EnSpm* | 0 | - | - | - | 1 | 0.3 | 0.0 | 1.0 |
|  | *hAT-Ac* | 1 | 3.6 | 0.0 | 10.9 | 3 | 5.9 | 0.0 | 14.7 |
|  | *P* | 0 | - | - | - | 1 | 9.0 | 0.0 | 27.1 |
|  | *PIF-ISL2EU* | 1 | 10.1 | 0.0 | 30.3 | 1 | 3.3 | 0.0 | 10.0 |
| LINE | *I* | 0 | - | - | - | 1 | 4.4 | 0.0 | 13.1 |
|  | *Penelope* | 1 | 11.0 | 0.0 | 32.9 | 0 | - | - | - |
| LTR | *DIRS* | 0 | - | - | - | 2 | 10.2 | 0.0 | 27.2 |
|  | *Gypsy* | 59 | 7.9 | 3.5 | 13.4 | 11 | 6.1 | 1.0 | 13.1 |
|  | *Pao* | 4 | 0.8 | 0.1 | 1.8 | 8 | 7.2 | 1.2 | 15.6 |

\*Confidence intervals estimated using by bootstrapping across MA lines 10000 times.

Averaging across rates for all TE families with non-zero copy numbers in all genotypes, there is evidence for highly variable gain and loss biases across genotypes (Figure 3A) and a binomial mixed effects model shows populations differ in their overall gain (Figure 3B, filled columns; χ^2^ = 6.22, df = 2, P =0.0446; Finland > Germany > Israel) and loss rates (Figure 3B, empty columns; χ^2^ = 12.07, df = 2, P =0.0024; Germany > Israel > Finland; Table S9A). In addition to the gain and loss rates, we also calculated a net and total mutation rate for each genotype (Table S9B) and for each active TE family (Table S10). These rates range from negative (e.g., *P* elements in genotype IB are decreasing at a rate of -7.44 x 10^-4^ per copy per generation) to positive (e.g., Penelope elements increasing at a rate of 9.06 x 10^-4^ /copy/gen in IC), and can even vary among genotypes for the same TE family (e.g., -2.22 and 4.06 x 10^-4^ /copy/gen for Gypsy in GA and FC, respectively). In comparison to the MA lines, the rates of gain and loss observed in the extant control lines (maintained in large populations in parallel to the MA lines, so selection was *not* minimized) were much lower and thus less variable among genotypes and TE families (Table 2; Table S10A and B). Differences in the number of events observed in the MA lines (where selection was minimized) compared to extant control lines (ECs) raised in large population sizes (where competition for resources means selection can occur) reveals the deleterious effects of TE activity, especially for gains. While the number of losses observed in the EC lines was less than a third of those seen in the MA lines, the reduction in the number of gains observed was much larger (2 [EC] versus 67 [MA] events; Table 2 and Tables S11A and B).

**Figure 3.**
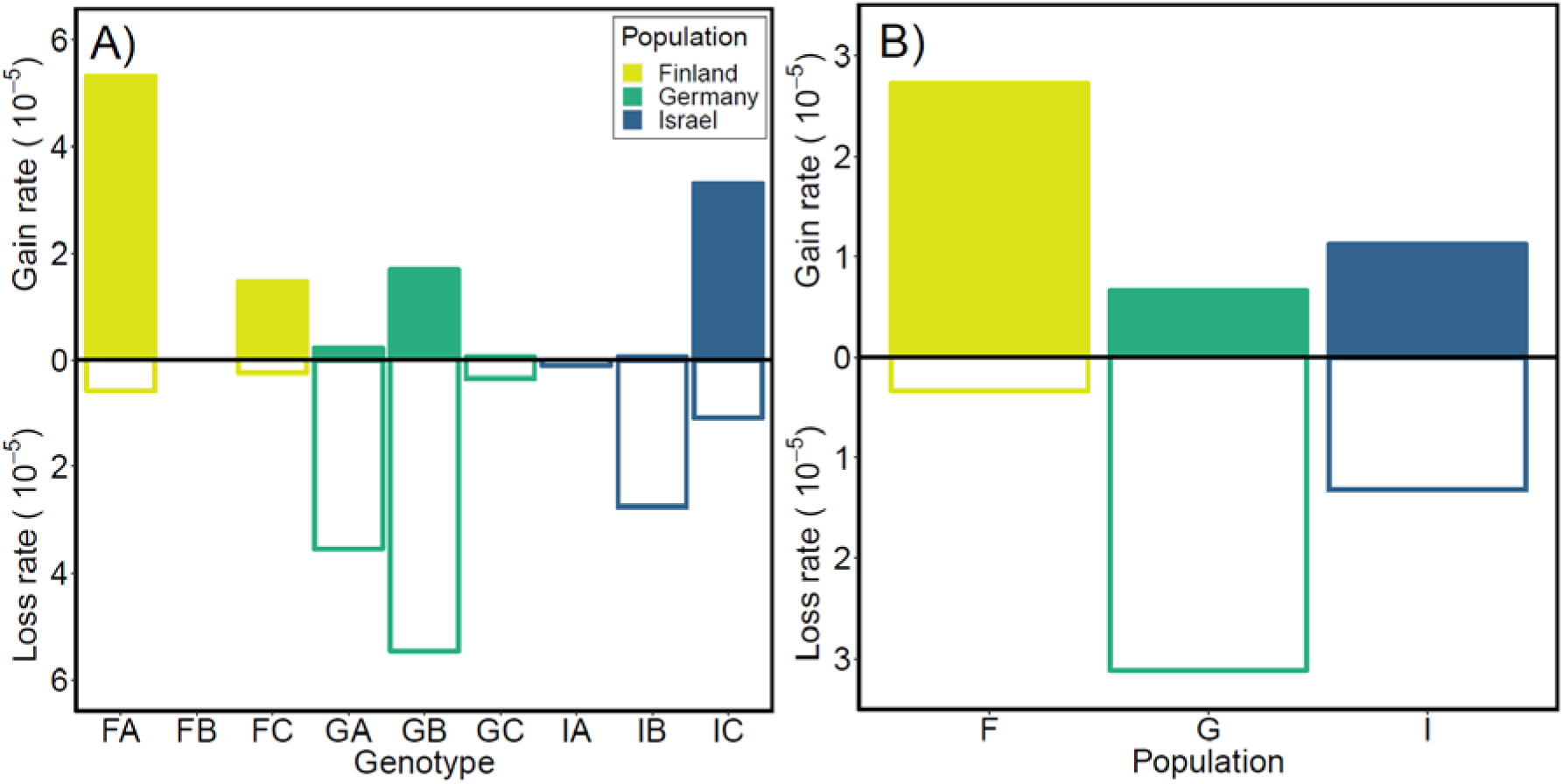
Gain and loss rates (per copy per generation) for each (A) genotype and each (B) population of *D. magna* averaged across all TE families. Mean rates for MA lines from Finland, Germany and Israel represented in gold, green and blue, respectively.

It is important to note, our TE mutation rate estimates represent a lower bound. They are based on the 67 TE gain and 28 TE loss events observed in 10 of 29 TE families in MA lines after filtering (see Methods). Unless otherwise noted, our analyses focus on only those TEs that could be classified as belonging to one of the five major groups of TEs. We calculated overall rates of loss and gain for all TEs (classified and unknown) identified using RepeatModeler, as well (see Supplemental Results and Table S12A and B). To gauge our sensitivity, we used simulations to estimate the false positive and false negative rate (FPR and FNR) for the four cases of TE events (Figure 1). FPRs were relatively low (< 0.013) for all four types of mutations (see Supplemental Results; Table S13A), and neither FNRs and FPRs vary depending on TE length (Table S13B). Notably, the fact that the four cases of events are not equally likely (most gains were novel (0 → 1 [n = 62/67]) and most losses were at previously heterozygous sites (1 → 0 [n = 26/28]) is revealing about what proximal mechanisms explain the bulk of TE proliferation and loss (see Discussion).

### Correlations with TE mutation rates

Overall, TE mutation rates in *D. magna* vary intraspecifically a great deal (Figure 3) mirroring the high levels of intraspecific variation observed in other mutation rate estimates for this species (see Ho et al. 2019, 2020). In terms of frequency per site, TE mutations are intermediate among the other types of mutation examined so far in *D. magna* (i.e., microsatellite mutation rates are much higher (10^-2^) and nuclear and mtDNA base substitution rates are much lower (10^-8^ and 10^-7^, respectively), on a per site per generation basis). Our population-specific means (Figure 3B) show that, in addition to high levels of variation among genotypes, populations differ in their overall rates of TE gain and loss, with Finland exhibiting a high net rate of loss, Germany exhibiting a high net rate of gain, and Israel exhibiting a low net rate because gains and losses (while still quite prevalent in 2 out of the 3 genotypes assayed) occur at nearly equal frequency. Because TE mutations can result in relatively large gains and losses in terms of the number of nucleotides added or lost per event, we looked at the relationship between rates of TE gain and loss (and net rates) and the proportion of the genome that is TEs in each genotype and found no correlation (Table S14), other than the fact that, as expected, we observe more events in higher copy number families as expected (Figure S6).

Comparing TE mutation rate estimates measured for each genotype to estimates for other types of mutation reveals no strong correlations with nuclear base substitution mutation rates (Figure 4A) or microsatellite rates (Figure 4B; Table S15). There is a correlation between TE mutation rates and gene conversion rates, when only the rates for TE events that are likely to be caused by gene conversion events are plotted (Figure 4C). This is expected, but it is important to note that the correlation here (ρ = 0.83, t_7_ = 3.91, P = 0.0058) appears to be driven largely by one genotype with high estimates for both rates (GB). Despite the lack of a consistent, linear relationship between TE mutation rates and other types of mutation rates across all 9 genotypes, it is clear that certain categories of mutation are consistently more frequent in Finland (TE gains are higher, microsatellite deletions are more common, and rates of base substitution in the nuclear genome are highest; Ho et al. 2019, 2020). Similarly, genotypes collected from Israel show consistently low relative rates of TE loss and gain, low rates of microsatellite mutation, and low rates of nuclear base substitution mutation rates. This suite of elevated mutation rates is reflected in the accumulation of higher estimates of overall TE loads long-term (6.6% using repeat masking) and a larger overall genome size (121 Mb average) for genotypes from Finland compared to Germany and Israel (Table S3). Looking at the largest TE family in *D. magna*, Gypsy, individually is illustrative, as it shows a close correspondence between genotype-specific mutation rates (high gain rates in Finland and high loss rates in Germany; Figure 5A), patterns of insertion site polymorphism (an excess of singletons in Finland and a paucity of population-specific insertions in Germany; Figure 5B), and the relative abundance of this TE family in the genome (high in Finland and low in Germany; Figure 5C and Table S15).

**Figure 4.**
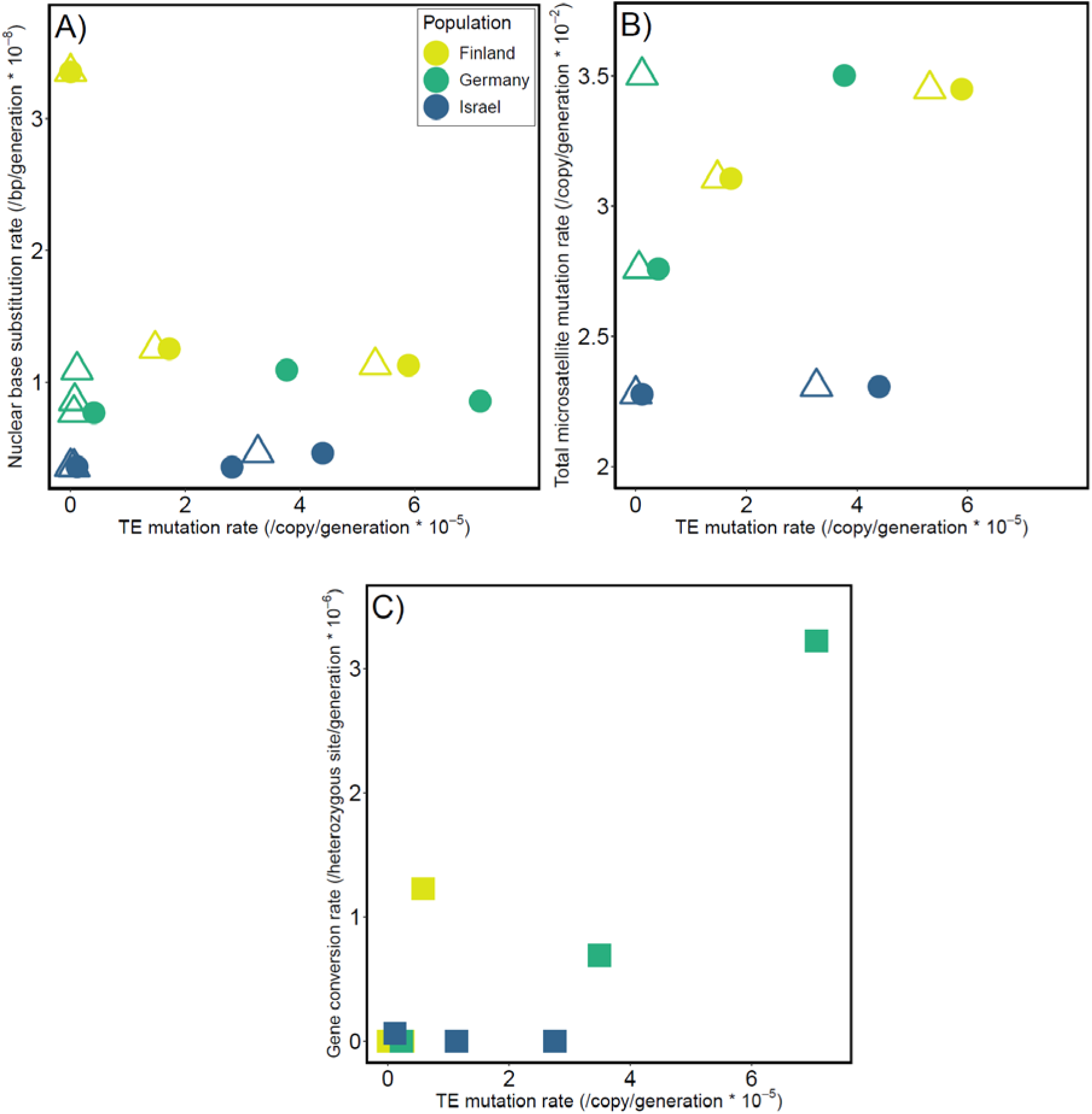
The relationship between TE mutation rates (per copy per generation) and (A) nuclear base substitution rates (per bp per generation), (B) absolute microsatellite mutation rates (per copy per generation), and (C) gene conversion rates (per heterozygous site per generation). TE rates (either the sum of all TE gains and losses (circles) or 0-->1 TE gain rate (triangles)) are plotted in gold (Finland), green (Germany), or blue (Israel). Microsatellite mutation rates are only available for 6 of the 9 genotypes (2 per population [Finland, Germany, and Israel]) and nuclear gene conversion rates are only plotted against events that could be caused by gene conversion (the sum of 1-->0 TE losses and 1-->2 TE gain rates; shown as squares).

**Figure 5.**
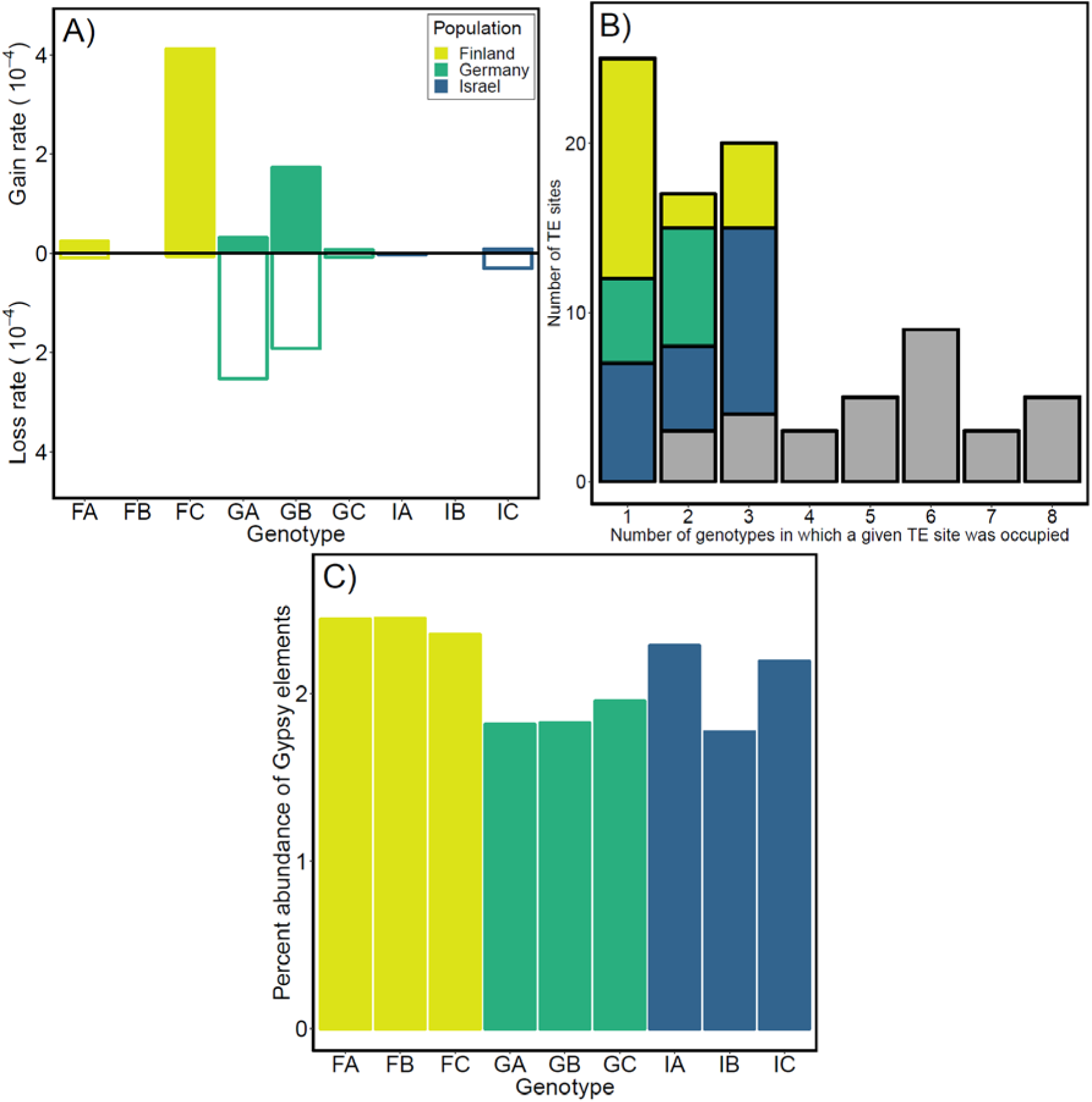
(A) Mean gain and loss rates (per copy per generation) of TEs from the Gypsy family for each genotype. (B) Count of polymorphic Gypsy sites separated by the number of SC lines (x) that contained Gypsy elements. Gypsy elements are singletons (genotype specific) when x=1 or are population specific when x=2 and x=3 (color indicates the populations the TEs are specific to). Grey represents sites where Gypsy elements are shared across populations. The reference genome used for this analysis was FASC. (C) Percent abundance of Gypsy elements in each of the assemblies estimated using RepeatMasker (Smit et al. 2013).

## Discussion

Our analyses of TE profiles among lineages over long-time scales and in real-time aims to (1) quantify the levels of intra- and interspecific variation in the TE content of the genome along a latitudinal gradient and among congeners, (2) estimate and compare TE mutation rates across genotypes and populations within a species, and (3) assess if long-term patterns of accrual can simply be explained by short-term mutational dynamics, or if they are heavily influenced by differences in the abiotic, biotic, or population-genetic environment among populations. This study represents the first such comparison on a large spatial scale, that includes intra- and interspecific comparisons, and provides the opportunity to directly compare TE mutation rates to those calculated for other categories of mutation in the same starting genotypes.

In order to pursue our goals, a number of challenges related to analyzing TEs and repetitive sequences in whole genome sequence datasets were addressed. First, characterizing the TE content in the genome is a major challenge (Arkhipova 2017). In fact, improvements in sequencing technologies, assembly algorithms, repeat-finding software and pipelines, and a deepening knowledge of TEs in other organisms over time means that the annotation of the TE profile in the genome of a given species is likely to improve over time which means it can be hard to standardize or compare across studies (Hoen et al. 2015). Our characterization of the TE content in *D. magna*, for example, included the identification of numerous TEs that cannot currently be classified into any of the major known categories of mobile elements (Piégu et al. 2015), and therefore could not be included in the calculations of family-specific rates (but see Tables S12A and B for overall rates including these TEs). In addition to finding and characterizing TEs in order to construct a library of elements, the most common method of quantifying repeat content in the genome (RepeatMasker; Smit et al. 2013) depends on quality of the genome assembly used, raising concerns about the accuracy of these commonly-used methods compared to less assembly-biased approaches where reads are mapped to the repeat library directly to estimate content (see Supplementary Results). Lastly, our method for measuring TE mutation rates (TEFLoN; Adrion et al. 2017) depends on being able to map reads that span gain and loss events, meaning read depth or length can alter the false positive and false negative rates. In short, while we believe the analyses we have conducted represent a thorough and consistent characterization of the repeat content and the TE mutation rates in each genotype, we present these data as lower-bound estimates as our ability to detect TEs in the genomes of emerging model organisms is likely to continue to improve with technological and bioinformatic advances (Table S13A and B).

Previous work on TEs in *Daphnia* focused mainly on differences between sexual and asexual reproduction given that cyclical parthenogenesis (asexual reproduction with occasional bouts of sex) makes this an excellent system for exploring the role of sex in the proliferation of TEs (e.g., Schaack et al. 2010a,b and Jiang et al. 2017). Unlike *Drosophila*, in which most TE rate studies in animals have been conducted (reviewed in Mérel et al. 2020), *Daphnia* can be propagated in the lab exclusively via asexually-produced clonal offspring allowing us to investigate the mutational processes associated with TE gain and loss, without the complicating influence of sex. The outcomes of these past studies have painted a complex picture, with some elements exhibiting different patterns of proliferation among sexuals and asexuals in some cases (e.g., Valizadeh and Crease 2008), but not all (Bast et al. 2016). Here, instead of contrasting sexuals and asexuals, we take advantage of our ability to subtract the influence of meiosis and syngamy on TE dynamics, and compare the mobile DNA profiles across 9 starting genotypes of *D. magna* originally collected from Finland, Germany, and Israel (Figure 2A) where mean temperatures, temperature ranges, light exposure, and the frequency of dry down differ (Table S1).

Overall, TE profiles were similar among genotypes from the three populations (Figure 2B) in terms of the abundance and diversity of TEs present. Across genotypes, elements from the Gypsy superfamily of LTRs (Class 1) are the most numerous, as has been reported in the congener, *D. pulex* (Rho et al. 2010), which has many more TEs overall than *D. magna* (Table 1), regardless of method used. Despite these similarities in patterns of accrual overall, insertion site polymorphism (differences among individuals in terms of which specific sites are occupied by TEs of a given family) distinguishes the three populations clearly (Figure 2C). Those distinctions are traceable to the high number of singletons found in genotypes from Finland, in contrast with the dearth of singletons (or even population-specific insertions) found in genotypes from Germany (Figure 2D). Finnish genotypes experience freezing temperatures and yearly dry downs, whereas German genotypes experience only freezing temperatures and genotypes from Israel experience only seasonal dry downs (Lange et al. 2015), potentially influencing the effective population size locally and thus the ability of selection to efficiently purge mutations that increase the mutation rate. Thus, the abiotic, biotic, and population genetic environments experienced by animals originating from these three different populations are distinctive, we were curious if mutation rates alone could explain these differences.

To estimate TE mutation rates, we performed a set of multi-year MA experiments initiated from each of the 9 starting genotypes of *D. magna* for which we characterized the TE content in the genome. After sequencing the whole genomes of descendant individuals from the MA lines and the extant control lines (lines not subject to population bottlenecks each generation, but instead maintained in large population sizes), we were able to count the number of gains and losses of TEs observed and calculate rates. Based on observations of over 100 gains and losses over an average of 12 generations of mutation accumulation across lines, we calculated overall rates of gain and loss (Table 2), as well as genotype- and population-specific (Figure 3) and rates for each superfamily/family of TEs (Table 3). Overall, rates of gain and loss in *D. magna* are similar (1.4 and 1.7 x 10^-5^ per copy per generation, respectively), but they vary a great deal among genotypes and populations (Figure 3 A and B). The majority of the gains observed are novel gains (1→0 gains; Figure 1) most likely resulting from insertions of TEs either excised from elsewhere in the genome (in the case of cut-and-paste elements) or retrotransposed (in the case of Class I elements, such as Gypsy). The majority of loss events were also observed at positions that were initially heterozygous (1 → 0), as would be expected given that previously homozygous loci may reconstitute TEs that excise or get deleted via homolog-dependent DNA repair (Engels et al. 1990). In *Drosophila*, a recent genome-wide assay of TE mutation rates showed insertions also far outnumber deletions, but overall all the per copy per generation rates are much lower and differ significantly, with insertions higher (∼10^-9^) than deletion rates (∼10^-10^; Adrion et al. 2017).

Little is known about intraspecific variation in TE mutation rates in animals, even though there have been several large-scale studies of their polymorphism (e.g., Petrov et al. 2010, Laricchia et al. 2017, and Lerat et al. 2019). Among *D. magna* genotypes, rates ranged from a high gain bias in a genotype from Finland (FA; 5.3 x 10^-5^ per copy per generation) to a deletion bias in one from Germany (GB; -5.5 x 10^-5^ per copy per generation; Table S9B), with the highest number of events by far occurring in a single genotype (FC; Table S9B) in a single family (Gypsy; Table S10). Looking across families of TEs, populations are distinct in their rates across TE families, with Finland exhibiting higher rates of gain overall, Germany exhibiting high rates of loss overall, and Israel exhibiting gains and losses with almost equal frequency resulting in the lowest net rates overall (Figure 3B; Tables S9A and S9B). Population-level variation may explain the distinctive patterns of insertion site polymorphism observed in the comparison of the TE profiles of the 9 starting genotypes from which the MA experiments were initiated. Furthermore, high levels of population-specific TE mutation rates and spectra may explain how two congeners (*D. magna* and *D. pulex)* with similar morphologies, life histories, and global distributions evolved substantially different TE fractions in their genomes overall.

Generally, rates were much higher in MA lines than in extant control lines, suggesting that TE mutational events, especially gains, are deleterious (Table 2). Although it has been posited that high copy number families might evolve to attenuate their rates of increase because of the deleterious effects of TE insertions (much like parasites evolve to lower virulence; Charlesworth and Charlesworth 1983), the relationship between per copy per generation rates of mutation and abundance in the genome observed here is weak overall (Figure S6C), with no clear downward trend even for rates of gain and relatively high rates being observed in the most abundant family, Gypsy (Table 3; Figure S6B). Even within a single family, rates of gain and loss can vary great among genotypes (e.g., Gypsy [Figure 5A]). This variation can be seen reflected in the patterns of polymorphism among genotypes (Figure 5B) and the overall abundance of that TE family in the genome (Figure 5C). Specifically, high rates of gain in Finnish genotypes could explain the excess of Gypsy singletons in these genotypes and the higher proportion of the genome that Gypsy elements occupy. Conversely, German genotypes had higher rates of loss for the family of TEs, have fewer singletons and no population-specific Gypsy insertions, and the lowest proportions of Gypsy in their genomes. It is worth noting that we observed the greatest number of events in the Gypsy superfamily, but on a per copy basis other families were also highly active. This is consistent with the idea that the large Gypsy family may be comprised of a mixture of active and inactive elements, as has been found in other systems (e.g., Lerat et al. 2003).

Previously, we reported estimates of microsatellite and base substitution mutation rates (bsMRs) and gene conversion rates for *D. magna* using the same genotypes assayed here (Ho et al. 2019, 2020). Microsatellites are known to be highly mutable (reviewed in Ellegren 2004)) and the average genome-wide rates of mutation at these loci in *D. magna* are several orders of magnitude higher (∼10^-2^) than the TE mutation rates we estimate here (∼10^-5^). The base substitution mutation rates we reported for *D. magna* were the highest and most variable direct estimates reported in animals using an MA framework (∼10^-9^ for the nuclear bsMR rates; Ho et al. 2020), but are, on average, several orders of magnitude lower than the overall TE mutation rates we report here. This highlights the fact that, while not frequently measured or modeled, rates of mutation other than base substitution rates are critical to quantify and understand, as they may be much greater sources of genetic variation, both in terms of frequency and the size of the mutations that occur, than the simple substitution of one nucleotide for another (Press et al. 2019). While bsMRs will likely remain the most commonly measured mutation rate, our prediction that there would be little to no correlation between those rate estimates with TE mutation rates bore out (Figure 4A and B), even though they do positively correlate with microsatellite rates (Ho et al. 2020). This may not be surprising, given that the mechanisms underlying TE insertions and excisions (e.g., the availability of functional transposase) and the plurality of mechanisms that can contribute to TE mutation rates (e.g., duplications, deletions, and gene conversion events) are not only numerous, but in large part distinctive from the main mechanisms thought to produce base substitutions (faulty DNA polymerase or DNA repair) or microsatellite expansions (DNA polymerase slippage). Nevertheless, evolutionary theory tends to aggregate all mutation types and models often depend on “the” mutation rate, which all too often is based on base substitution mutation rates which, even when derived from an MA experimental framework where selection is minimized, might not provide an accurate measure of the rate at which genetic variation is being introduced.

Despite the lack of correlations among mutation rates across genotypes, it is interesting to note that rates of gene conversion (from Ho et al. 2020) do correlate with the rates of TE events when rate averages are based on only those events that can be produced by gene conversion events (Figure 1; Figure 4C; and Table S14). Generally, rates do not correlate with the abundance of TEs in the genome overall (Table S14), meaning that the overall bias towards gains in Finland and towards losses in Germany does not appear to have a significant effect on genome size. While, using the commonly used repeat masking approach to estimating TE content, it appears that *D. magna* have fewer TEs (proportionally) than *D. pulex*, this could be an artifact of having a repeat-rich genome. Using the repeat mapping approach, the proportion of the genome estimated to be TEs in *D. magna* is 16%, but this number is difficult to compare across studies without access to the unfiltered reads. The accumulation of TEs has been offered as a solution to the C-value paradox (Biscotti et al. 2019), but it remains an unresolved question whether TE content explains the larger genome size of *D. magna* relative to *D. pulex* based on cytological methods (Jalal et al. 2013). These congeners exhibit a number of unexpected differences in their mutation profiles, including major differences in base substitution mutation rates (*D. magna* >> *D. pulex*; Ho et al. 2020) and microsatellite mutation rates (*D. magna* >> *D. pulex*; Ho et al. 2019). Future work on TE mutation rates in *D. pulex* and other species will reveal useful insight into how species differ in these important evolutionary parameters across near and distant branches of the tree of life (e.g., Sayres et al. 2011).

To determine if the TE mutation rates reported here include some genotypes with rates at the upper end of the range for the species, or if this sampling represents a high mean estimate compared to other eukaryotes requires additional data. Few direct estimates of TE mutation rates have been published outside of *Drosophila* (e.g., Nuzhdin and Mackay 1995), human (e.g., Feusier et al. 2019), and a few species of plants (e.g., Quadrana et al. 2019), and the levels of rate variation must be established in order to pursue deeper inquiries into the causes and consequences of such variation. In our analysis, the high mean rates of gain observed in Finnish genotypes could explain the excess of singletons among genotypes and the higher proportion of TEs found in these genotypes compared to those from Germany and Israel (as illustrated for Gypsy elements in Figure 5). That being said, the differences in insertion site polymorphism and TE abundance between populations should be interpreted with caution, given we only have three genotypes per population. Biotic or abiotic conditions, or historical population genetic constraints experienced by genotypes in Finland, but not the other genotypes, could also dictate the selection regime shaping the evolution of the mutation rate in these three populations. For example, periodic dry-downs and freezing temperatures could both impact DNA molecules physically, affecting the mutation rate, or could cause frequent genetic bottlenecks in Finnish populations, reducing the local effective population enough to weaken the efficiency with which natural selection can remove mutator alleles, thereby increasing the mutation rate.

While there is no positive linear correlation among mutation rates for different types of mutations (TEs, base substitutions, and microsatellites) across all 9 genotypes, it is interesting to note that, in a mutation accumulation experiment, genotypes from Finland have the highest rates of TE gain (and gains are more deleterious than losses), the highest rates of microsatellite deletions (Ho et al. 2019), and the highest rates of base substitution among the three populations assayed (Ho et al. 2020). These commonalities among our direct estimates based on rearing animals in a common laboratory environment point to population genetic constraints, rather than mutagens in the atmosphere, as an explanation for higher rates of deleterious mutation in the Finnish genotypes. In contrast, genotypes from Israel consistently exhibit the lowest net rates of TE mutation, microsatellite mutation, and base substitutions of the three populations assayed. Early puzzles about mutation rate variation, for example the divergence of mitochondrial base substitution mutation rates relative to those in the nucleus between plants and animals (Lynch et al. 2006), remain unsolved (Havird and Sloan 2016) and the investigation into other mutation types will undoubtedly reveal more questions than answer, at first. Future work aimed at understanding the causes and consequences of mutation rate variation within populations and species, the heritability and evolvability of mutation rates for different types of mutation, and the significance of the mobilome for generating genetic variation are necessary to improve our understanding of how mutation rates evolve over time and space.

## Methods

### Study System

The *D. magna* genotypes used in this experiment were collected along a latitudinal gradient that captures a range of environmental variation, including temperature and photoperiod (Table S1), and which can result in fluctuating habitat sizes (see Yampolsky et al. 2014). Three genotypes from each of three populations (Finland, Germany, and Israel) were used to initiate lab stocks from which the starting controls (SCs) were picked, from which the mutation accumulation (MA) lines and extant controls (ECs) were then propagated. For each of the 9 genotypes, the SCs were frozen tissue from the individuals whose descendants were used to initiate the MA lines and ECs for that genotype, for which tissue was frozen after the mutation accumulation period (the average number of generations across MA lines was 12 and the experiment ran for approximately 30 months in total).

The MA lines from each genotype were maintained as single individuals in 250 mL beakers containing 175-200 mL of Aechener Daphnien Medium (ADaM; Klüttgen et al. 1994) under a constant photoperiod (16L:8D) and temperature (18° C), and fed the unicellular green alga *Scenedesmus obliquus* (2-3 times per week *ad libitum*). The ECs for each genotype were maintained in large population sizes in 3.5 L jars in the same medium, environmental conditions, and were also fed *ad libitum*. While selection is permitted to act in the large population ECs, the single-progeny descent used to propagate the MA lines minimizes selection and thus allows for mutation accumulation. At the end of the mutation accumulation period, the SCs, MA lines, and ECs were all sequenced to assess the TE content in the original genotypes (SCs), to assess TE mutation rates (MA lines), and to compare to lab-reared lines where selection is not minimized (ECs). The MA experiment used here has been described previously (Ho et al. 2019, 2020).

### DNA Extraction and Sequencing

Five asexually-produced clonal individuals from each SC line, all derived MA lines, and the extant control lines were flash frozen for DNA extractions. DNA was extracted using the Zymo Quick-DNA Universal Solid Tissue Prep Kit (No. D4069) following the manufacturer’s protocol (DNA from a few samples was also extracted with the Qiagen DNeasy Blood and Tissue Kit, No. 69504). DNA quality was assessed by electrophoresis on 3% agarose gels and DNA concentration was determined by dsDNA HS Qubit Assay (Molecular Probes by Life Technologies, No. Q32851). The Center for Genome Research and Biocomputing at Oregon State University generated 94 Wafergen DNA 150bp paired-end libraries using the Biosystems Apollo 324 NGS library prep system. Quality was assessed using a Bioanalyzer 2100 (Agilent Technologies, No. G2939BA) and libraries were pooled based on qPCR concentrations across 16 lanes (2 runs). Libraries were sequenced on an Illumina HiSeq 3000 (150 bp PE reads) with an average insert size of ∼380bp to generate approximately 50x coverage genome-wide for each sample. See Table S2A and B for genome assembly statistics for each whole genome sequence (WGS) produced.

### TE Characterization

To reduce reference bias, we searched for TEs using reference-guided assemblies for each of the 9 genotypes (using the WGS from each SC). Reads from each sample were processed by trimming adaptor sequences (k=23, ktrim=r, mink=4, hdist=1, tpe, tbo), merging overlapping pairs (vstrict=t), and quality filtering (qtrim = rl, trimq=20, minlen=50) with BBTools (Bushnell et al. 2017). First pass reference-guided *de novo* assemblies were performed with SPAdes (using the *trusted-contigs* option; Bankevich et al. 2012). The *D. magna* reference used to guide the assembly was provided by Peter Fields and Dieter Ebert (personal communication). To remove haplotigs potentially derived from assembling heterozygous regions into alternate alleles, we collapsed the filtered assemblies with *redundans* by implementing information from paired-end reads, merged reads, and the D. magna reference genome for scaffolding (Pryszcz and Gabaldón 2016). We then mapped the processed reads of each SC onto the collapsed assemblies with BWA-MEM v0.7.17 (Li and Durbin 2010). Contigs were removed if they possessed an average depth of coverage <5 or were shorter than 5 kb.

### Calculating TE Abundance and Diversity, Insertion Site Polymorphism and Mean Pairwise Divergence

A custom *D. magna* TE consensus library was created from a concatenated file of the assemblies from the 9 genotypes’ SCs using RepeatModeler v1.0.11 (Smit & Hubley 2008). Each assembly was then masked with our custom TE consensus library, using the slow search setting of RepeatMasker v4.1.0 (Smit et al. 2013). The output of RepeatMasker was used to determine the number, length, pairwise divergence, class and family of transposable elements in each assembly. For each assembly, we calculated the proportional abundance of each TE class or family by dividing the sum of their lengths by the total length of the assembly. We also used a read-mapping approach to calculate the TE fraction of the genome. For each assembly, we clustered elements in the TE library if they exhibited at least 98% nucleotide identity over their full length to a longer sequence in the library with the program cd-hit-est v4.8.1 (Fu et al. 2012, Li and Godzik 2006), yielding a non-redundant TE library containing full and partial TE copies. Elements with matches to sequences in the consensus library shorter than 150 bp, the average sequence read length, and identified as “Unknown” were excluded from this non-redundant TE library. Processed reads were then mapped to the non-redundant TE library with BWA-MEM v0.7.17 (Li and Durbin 2010). The number of bp in the genome covered by a particular TE copy was determined by summing the lengths of reads mapped to each TE sequence and normalizing to the median coverage of single-copy regions for each sample. The lengths of all constituent TE copies were summed to estimate the total number of bp in each genome and divided by the length of the assembly to obtain percent occupancy.

To estimate the insertion site polymorphism (i.e., the number of TEs at a particular site in a genome shared by multiple genotypes) we used TEFLoN v0.4 (Adrion et al. 2017). TEFLoN allows users to determine the presence/absence patterns of TEs among lineages, either among natural isolates (i.e., our 9 starting genotypes) or among MA lines (i.e., the MA lines derived from each starting genotype). ‘Presence’ reads are soft-clipped at a TE insertion site, or have one end of a pair mapped to a TE copy while the other maps to a unique position in the pseudo-reference, while ‘absence’ reads span an insertion site in the pseudo-reference. Pseudo-references are created by removing all matches to TEs in the consensus library identified with RepeatMasker v4.1.0 (Smit et al. 2013) that are >150 bp. Non-mergeable, quality-filtered paired end reads are mapped to pseudo-references for corresponding SCs with BWA-MEM v0.7.17 (Li and Durbin 2010), with soft-clipping for supplementary alignments (-Y). As a result, TEFLoN reveals the presence/absence status at each site (TE insertion polymorphism [TIP]), although it does not return the specific internal sequence that occupies/does not occupy scored sites.

We specified an empirically determined insert size standard deviation of 150 bp, and removed TEs identified within 5 bp of the end of any scaffold. To identify TIPs, we mapped reads from each of the 9 SCs to the pseudo-reference of a focal SC, and repeated this using all 9 SCs as the focal. We required that all occupied/unoccupied sites were supported by ≥ 20 presence + absence reads with mapping quality ≥30. A TE copy was considered present in a line if the frequency of the presence allele was ≥0.05 (altering this threshold did not substantially alter the distribution of TIPs identified using each reference). Sites were excluded if one or more SC lines failed to pass this filter. We considered a site to be polymorphic if one or more SC lines did not contain a copy of the TE. Genotype-specific TIPs were defined as sites where only one SC line contained the TE. Population-specific TIPs were defined as sites where two or three SC lines from the same population contained the TE while the remaining SC lines did not contain the TE.

### TE Mutation Rate Estimation in MA Lines

We used TEFLoN v0.4 (Adrion et al. 2017) to identify active TEs in the MA lines. For each genotype, we first identified heterozygous sites in all lines (SC, MA, EC) where a site was supported by ≥ 20 reads (presence + absence) with mapping quality ≥ 30 and contained at least one presence and one absence read. For this set of heterozygous sites, we recorded the 10^th^ and 90^th^ percentile of the presence allele frequency (p), denoted as p_10_ and p_90,_ respectively. The median of p in each genotype was less than 0.5, suggesting that presence reads were less likely to map correctly. The mean p_10_ and p_90_ across genotypes were 0.11 and 0.76, respectively. Subsequently, for all sites where all lines (SC, MA and EC) were supported by ≥20 presence + absence reads, we assigned a particular line as homozygous for the presence allele if p > 0.95, homozygous for the absence allele if p < 0.05, heterozygous for the presence allele if p_10_≤ p≤p_90_ and otherwise as undefined. All sites containing one or more undefined lines were discarded from further analyses.

There are two types of TE gain mutations (0-->1 and 1-->2) and two types of TE loss mutations (2-->1 and 0-->1) that can be observed based on whether the ancestor (SC) was homozygous, heterozygous or lacked a TE (an “absence allele”) at a given site relative to the status in the descendant MA line (e.g., if the SC was heterozygous and experienced an gain, it would be classified as a 1-->2 gain event in the MA line; Figure 1). Our ability to detect these different events is not uniform, however, because the likelihood of detecting heterozygous and homozygous “presence” alleles is not simply doubled. Furthermore, because we observe that “presence alleles” in heterozygotes tend to have a frequency below 0.5, it is possible that a line we determined to be homozygous for absence alleles may actually be a heterozygote, but we only observed absence alleles because the presence allele had a higher mapping error rate. To help guard against these types of false positives, we applied an additional filter to the sites tentatively identified to have experienced a TE mutation event. For sites that tentatively experienced a 0-->1 gain or 2-->1 loss, we wanted to guard against the possibility that one or more of the homozygous non-focal lines were actually heterozygous but assigned homozygous due to mapping error. We estimated the frequency of presence and absence alleles for heterozygotes at the site using the frequency in the focal mutant heterozygous line. Given this frequency, we calculated the binomial probability that one or more of the homozygous lines were heterozygous and discarded the site if this probability was larger than 0.005. For sites that tentatively experience a 1-->2 gain or 1-->0 loss, we wanted to guard against the possibility that the focal mutant homozygous line was actually heterozygous but assigned homozygous due to mapping error. To estimate the frequency of presence and absence alleles for heterozygotes at the site, we took the average of frequencies across the heterozygous non-focal lines. Given this averaged frequency, we calculated the binomial probability that the homozygous focal line was heterozygous and discarded the site if this probability was larger than 0.005.

Family-specific mutations rates for each of the four mutation types were calculated using N_m_ / (N_SC_*G), where N_m_ represents that number of sites that experienced a particular mutation event, N_SC_ represents the initial copy number of that TE family in the SC line, and G represents the number of MA generations. Because many repeats cannot be characterized to the level of family, we also ran TEFLoN using the full library containing all known and unknown repeats to obtain the total number of gains and losses and calculate an overall (not family-specific) rate of gain and loss for all repeats found in the initial RepeatModeler run (Table S12A, S12B), however our downstream analyses focus on the rates calculated for those TEs that could be characterized.

### Estimating false positive and false negative rates

To estimate false positive rates (FPR) and false negative rates (FNR), we simulated the MA experiment and processed the data using TEFLoN v0.4 (Adrion et al. 2017) and our filtering pipeline (similar to Adrion et al. 2017, 2019). For each simulation, we generated 11 unique diploid individuals by simulating SNPs onto the largest contig of the FASC assembly (6.8 Mb) using pIRS v1.1.1 (Hu et al. 2012). The 11 individuals represented one ancestral line and 10 descendent (mutation accumulation + extant control) lines. We then inserted 50 of one type of TE mutation event: 0-->1 gain, 1-->0 loss, 1-->2 gain or 2-->1 loss. For each mutation, we randomly picked an existing TE on the contig with a minimum length of 400 bp and randomly inserted it back into the contig, making sure it did not overlap with other TEs. We included a target site duplication (TSD) flanking each insertion with a mean length of 5 bp drawn from a Poisson distribution. To simulate a 0-->1 gain, we inserted a heterozygous TE (i.e., on one homolog of the contig) into one of the descendent lines. To simulate a 2-->1 loss, we inserted a homozygous TE (i.e. insertion of both homologs) on the ancestral and all non-focal descendant lines and then inserted a heterozygous TE on the focal MA line. To simulate a 1-->2 gain, we inserted a heterozygous TE on the ancestral and all non-focal descendant lines and then inserted a homozygous TE on the focal MA line. Lastly, to simulate a 1-->0 loss, we inserted a heterozygous TE on the ancestral and all non-focal descendant lines and did not change the focal descendent line. Finally, we independently simulated pair-end reads for each of the 11 individuals with an average coverage of 50x using pIRS (Hu et al. 2012) and filtered for mutations in the same way as described above.

For each mutation type, we repeated the simulation four times. We also examined whether the minimum length of TEs affected FPR and FNR by repeating the whole process and setting the minimum length of TEs to 800 bp. In total, we simulated 400 of each mutation type. FPR for focal mutation type was estimated as FP / (FP + TN), where FP is the number of discovered TEs falsely inferred to be the focal mutation type, and TN is the number of existing TEs not identified as a mutation. FNR for the focal mutation type was estimated as FN / (FN + TP), where FN is the number of simulated mutations that were not identified and TP is the number of simulated mutations that were detected.

### Statistical Analyses

Statistical analyses were performed in R (R Core Team 2018). Family-specific TE mutation rates for a particular genotype was estimated by averaging across MA lines. Rates of a particular mutation type (0-->1 gain, 1-->2 gain, 1-->2 loss, 2-->1 loss) of a MA line was estimated by averaging that rate across all TE families. Rates of a particular mutation type for a genotype was estimated by averaging that rate across MA lines. Confidence intervals for mutations rates were estimated by bootstrapping across MA lines 10000 times. Rates for EC lines were estimated similarly to the MA lines. To test for differences in TE mutation rates, we used a binomial mixed-effects model on the number of gains (or losses) with population and genotype (nested within population) as the fixed and random effects, respectively. We used Pearson correlations to test for statistical association among rates for different types of mutations. Base substitution, gene conversion and microsatellite mutation rates were obtained from previous studies (Ho et al. 2019, 2020) on the same *D. magna* MA lines, although in some cases estimates are available for only 6 of the 9 genotypes. All code for data processing and analysis is available at https://github.com/EddieKHHo/DaphiaMagna_MA_TE and sequence data have been deposited at NCBI (PRJNA658680).

### Data Access

Whole genome sequence data have been deposited at NCBI (PRJNA658680), all code is available online (https://github.com/EddieKHHo/DaphiaMagna_MA_TE), and the TE library used will be made available as a supplemental data file upon acceptance.

## Acknowledgments

We would like to thank Maia J. Benner, Dana Howe, Dee Denver, Dieter Ebert, Peter Fields, and Jeremy Coate for technical assistance, resources/support, and helpful feedback. We would also like to acknowledge our funding sources: awards from the National Institute of General Medical Sciences of the National Institutes of Health (GM132861) and National Science Foundation (MCB-1150213) to SS.

## Supplemental Results

### Repeat masking versus read mapping

The disparity between methods for estimating TE abundance (repeat masking versus read mapping) is likely because more repetitive genomes are harder to assemble, and the estimation of repeat content using repeat masking depends on the quality of the assembly. In support of this notion, we observe a negative correlation between assembly size and proportion of the assembly containing TEs using a read mapping approach (ρ = -0.69, t_7_ = -2.49, P = 0.042), as would be expected if high repeat content impedes confident assembly. The abundance estimates based on the two methods differ considerably if you aggregate by TE type (ρ = -0.19, t_7_ = -0.49, *p* = 0.63), however when compared by individual TE family the methods correlate strongly (bp averaged across genotypes; ρ = 0.99, t_33_ = 34.1, *p* < 0.0001; Table S4).

### Rate estimates including unclassified (“unknown”) repeats

Our main analysis excluded a large number of interspersed repeats that were designated as “unknown” by Repeat Modeler v1.0.11 (Smit & Hubley 2008). To examine if these unknown repeats experienced mutational events, we ran our analysis again using a repeat library containing the known TEs from the main analysis and the unknown repeats. The analysis including the unknown repeats discovered 225 indels, which was greater than the 95 events in the main analysis (Table S12a). Of the 225 indels, 72 were associated with TEs and 153 with unknown repeats.

The effect of including unknown repeats in our analysis varied greatly among genotypes. While we observed a modest increase in mutations for GC, IA and IB, there was a larger increase of 29 and 57 events for FC and GB, respectively (Table S12B). Furthermore, we found 0 mutations for FB in our main analysis but our analysis with unknown repeats discovered 5 mutations. Interestingly, the distribution of gain and losses were relatively similar between TE and unknown repeats for FC and GB. In FC, we observed a bias for 0-->1 gains for both TE and unknown repeats while for GB the bias was for 1-->0 losses. Although these results suggest we may be underestimating the total count of TE mutations, the total rate of mutations was similar to our main analysis that excluded unknown repeats (Table S12b). In fact, the total rate was slightly lower for FC, GC and IB in the analysis that included the unknown repeats. This is reassuring because it suggests that some mutational properties of TEs can be approximated even if many TEs have not been fully characterized.

### Simulations to estimate false positive and false negative rates

We used simulations to estimate the false positive and false negative rate (FPR and FNR) for the four cases of events (Figure 1). Two categories of events (0-->1 gains and 2-->1 losses) have low FNRs (0.06 and 0.09, respectively), while 1-->2 gains and 1-->0 losses had higher FNRs (0.13 and 0.28, respectively; Table S13A). We determined that the higher FNRs for 1-->2 gains and 1-->0 losses were mainly due to the heterozygous TE sites not being detected by TEFLoN, possibly due to low coverage and/or mapping error errors for repetitive regions. False positive rates (FPR) were relatively low (< 0.013) for all four types of mutations (Table S12A), and neither FNRs and FPRs vary depending on TE length (Table S13B).

### Analysis of TE insertion site polymorphism for different reference assemblies

When searching for TE insertion site polymorphisms (TIPs), we had to choose a common reference assembly for TEFLoN to map reads against. Since we had nine reference assemblies, we ran this analysis nine times using each of the different assemblies as reference. Although the total number of TIPs found varied when using different assemblies (Table S5), the general patterns are robust across the nine analyses.

Firstly, we found no significant difference for the distribution of TIPs in each TE family (χ^2^ = 238.51, df = 240, P = 0.51; Table S5). The TE families Gypsy, Pao, hAT-Ac, Copia, I, and DIRs possessed the highest number of TIPs across all nine analyses. Secondly, principal components analysis was also able to cluster genotypes by their population of origin regardless of the reference assembly used (Figure S5). Lastly, neither the distribution for the frequency of TEs at TIPs (χ^2^ = 67.93, df = 72, P = 0.61; Figure S6) nor the proportion of singletons belonging to each population (χ^2^ = 8.8, df = 16, P = 0.92; Figure S1) were significantly different when using different reference assemblies. Across the nine analyses, 42-53% of TIPs were population-specific and 13-16% were singletons. In addition, Israel genotypes tend to possess the highest number of population-specific TEs, while Germany genotypes tend to possess the least. The relative lack of singletons TIPs in German genotypes helps explain why German genotypes tend to cluster less closely than Finnish and Israeli genotypes on our PCA plots (Figure S5).

---

**Figure S1.**
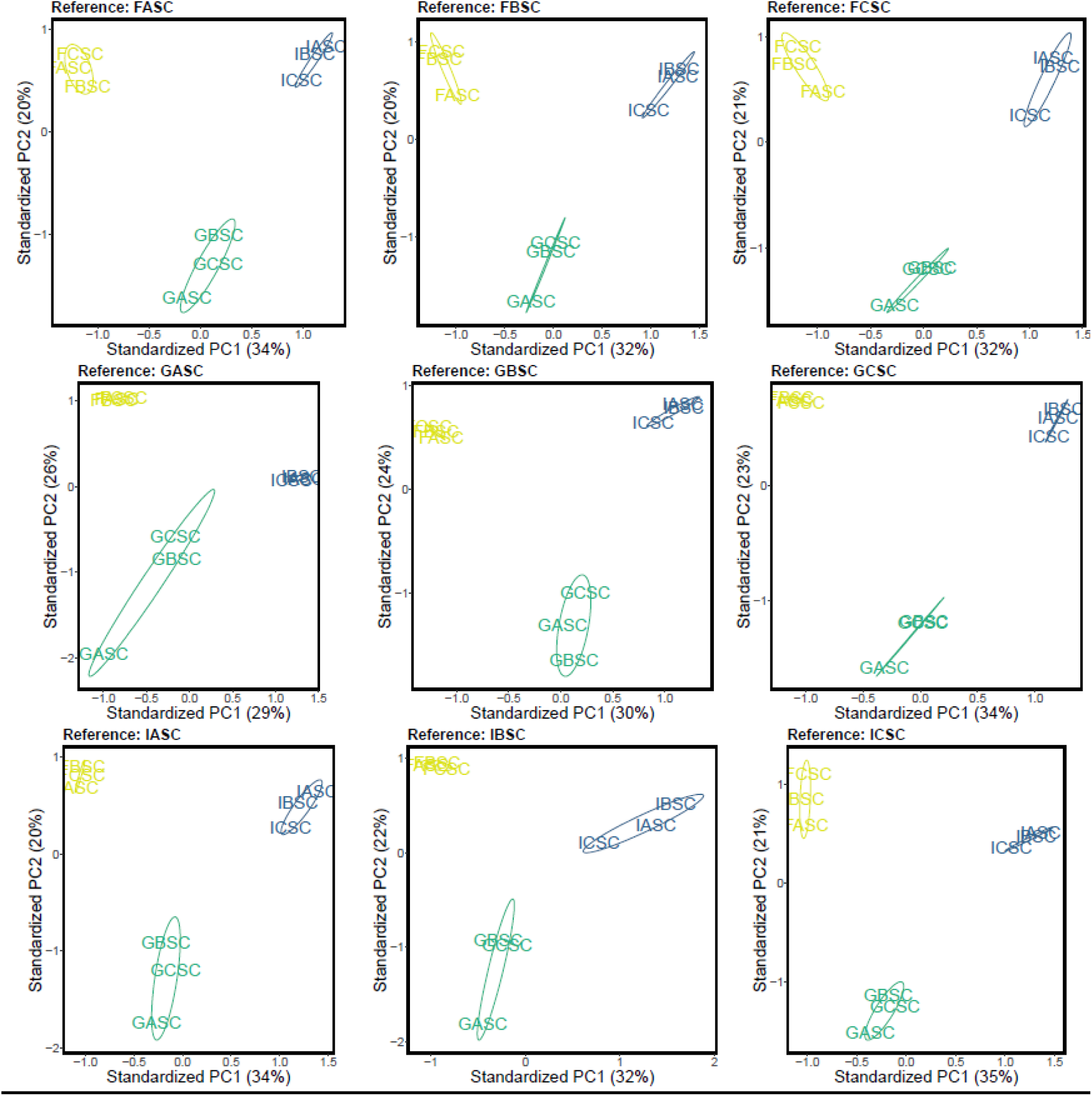
Principal Component Analysis based on the presence/absence of TEs when using each of the nine reference assemblies. Variance explained by principal components 1 and 2 are displayed on the axes. The reference assembly used is indicated on the top of each plot. Genotypes from Finland, Germany and Israel are colored in gold, green, blue, respectively.

**Figure S2.**
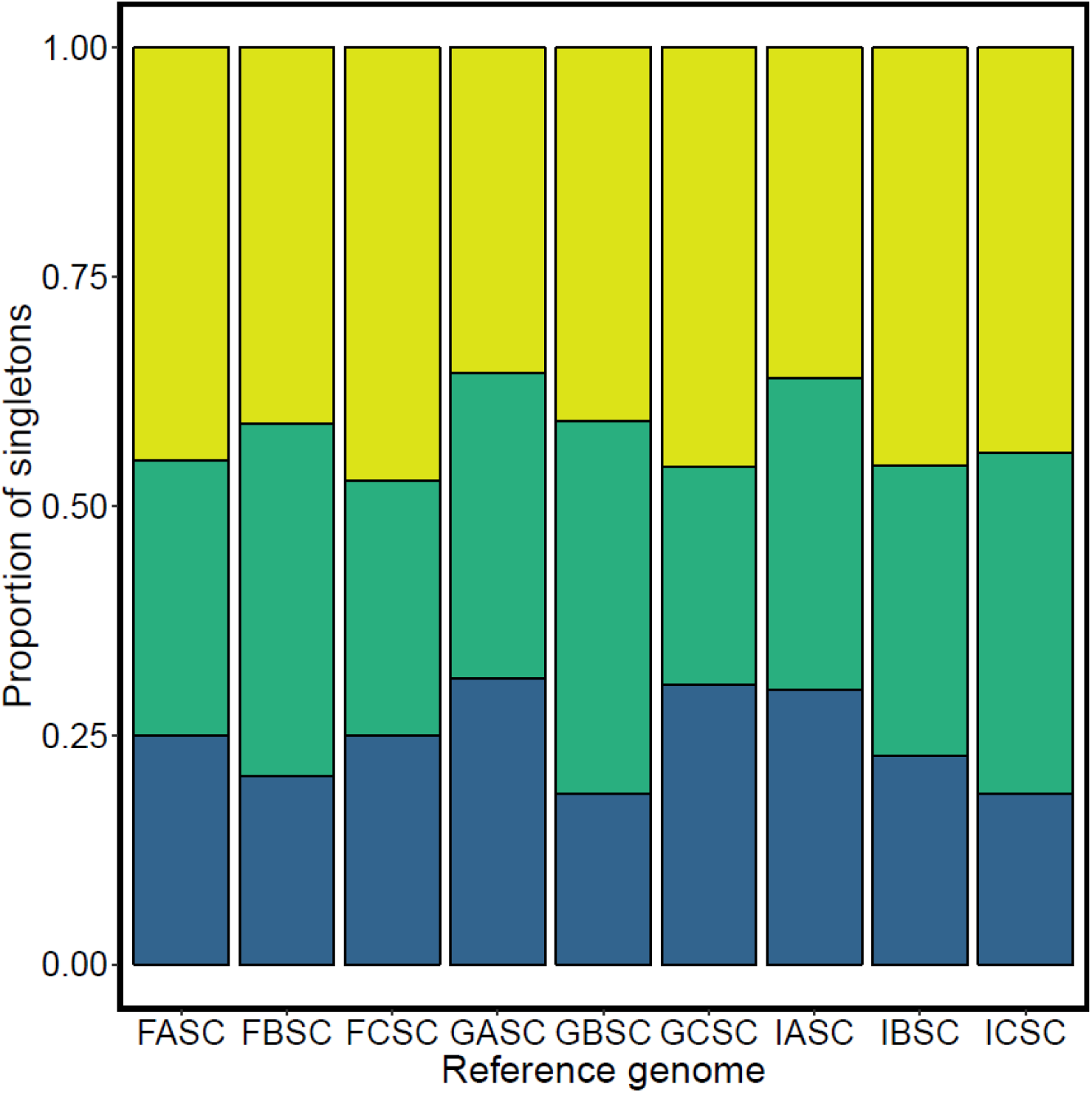
Proportion of singletons TEs in each population for analyses using different reference genomes. Gold, green and blue represents singletons specific to genotypes in Finland, Germany and Israel, respectively. The proportion of singletons belonging to each population was not significantly different when using different reference genomes (^2^ = 8.8, df = 16, P = 0.92).

**Figure S3.**
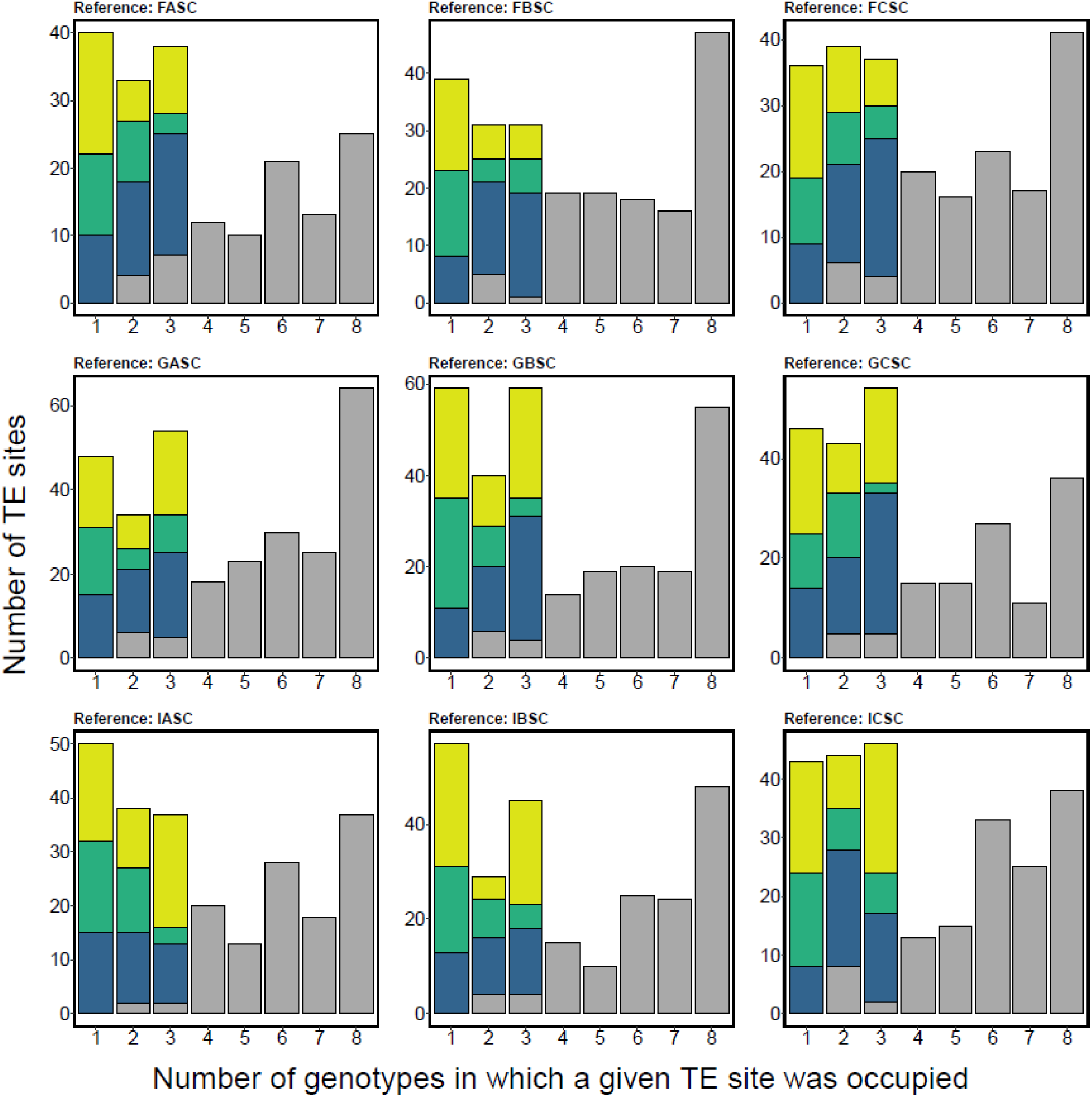
Number of polymorphic TE sites occupied across the 9 genotypes. The left bar represents the number of singletons (sites occupied in only one genotype) for each population (gold, green and blue for Finland, Germany and Israel, respectively). Colored portions of bars in x = 2 and x = 3 represent sites occupied in 2 and 3 genotypes, respectively, when from the same population. Grey portions of each bar represent the number of sites that were occupied in ≥2 genotypes that were not population-specific. The reference assembly used is indicated on the top of each plot.

**Figure S4A.**
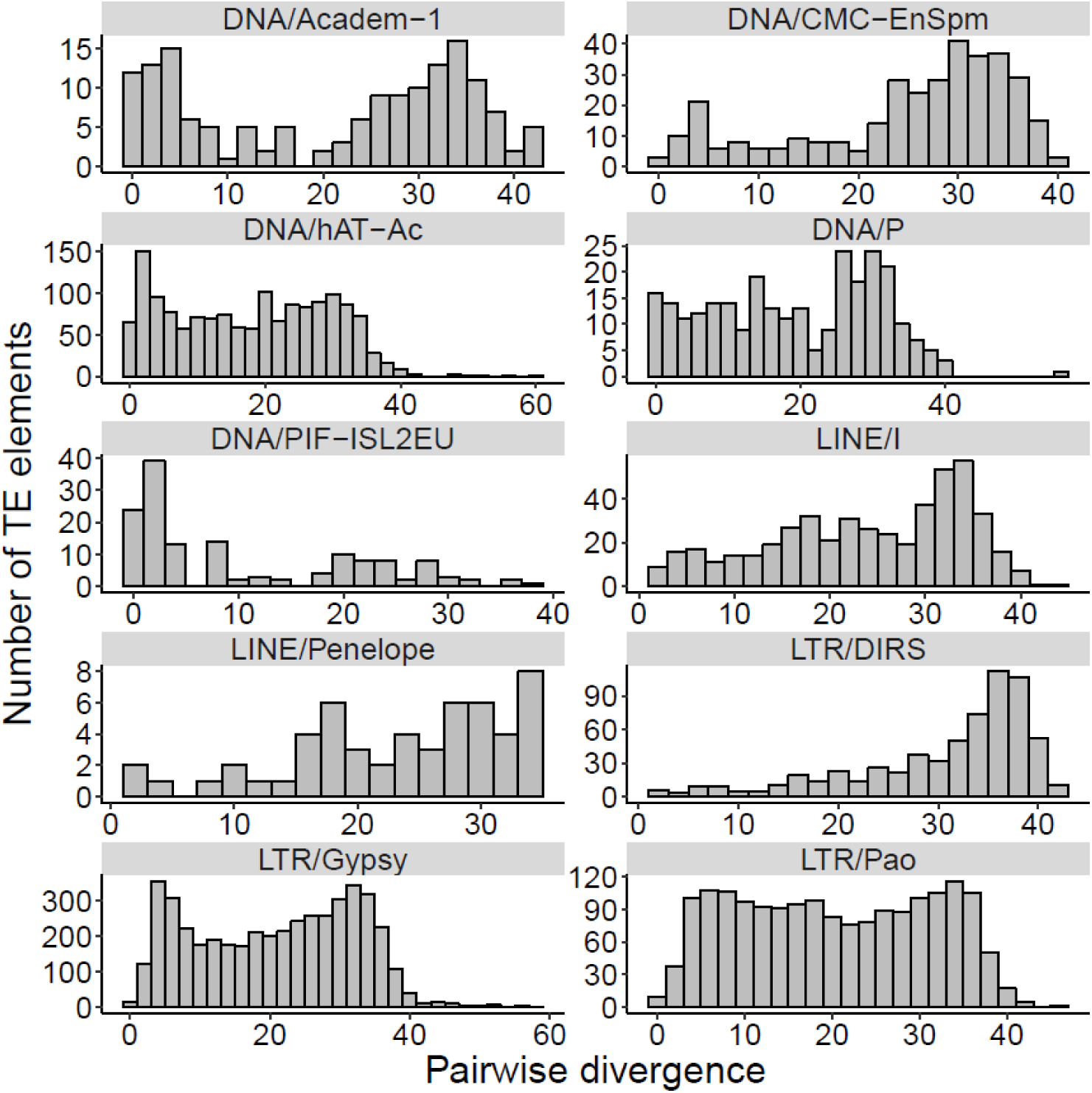
Pairwise divergence of TEs in the *D. magna* FASC assembly for TE families that are active within *D. magna* MA lines.

**Figure S4B.**
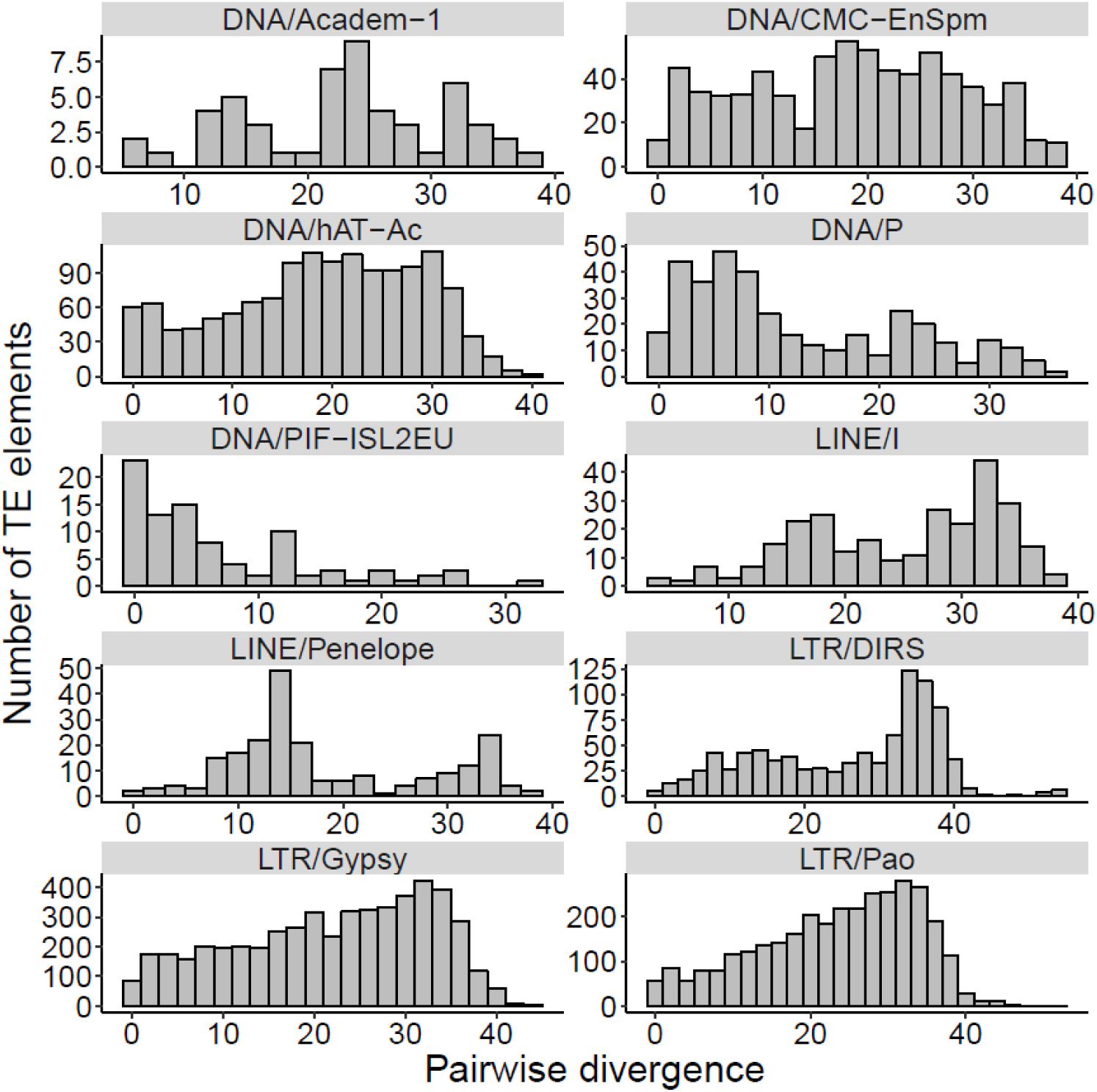
Pairwise divergence of TEs in the *D. pulex* reference genome (PA42) for families that are active within *D. magna* MA lines.

**Figure S4C.**
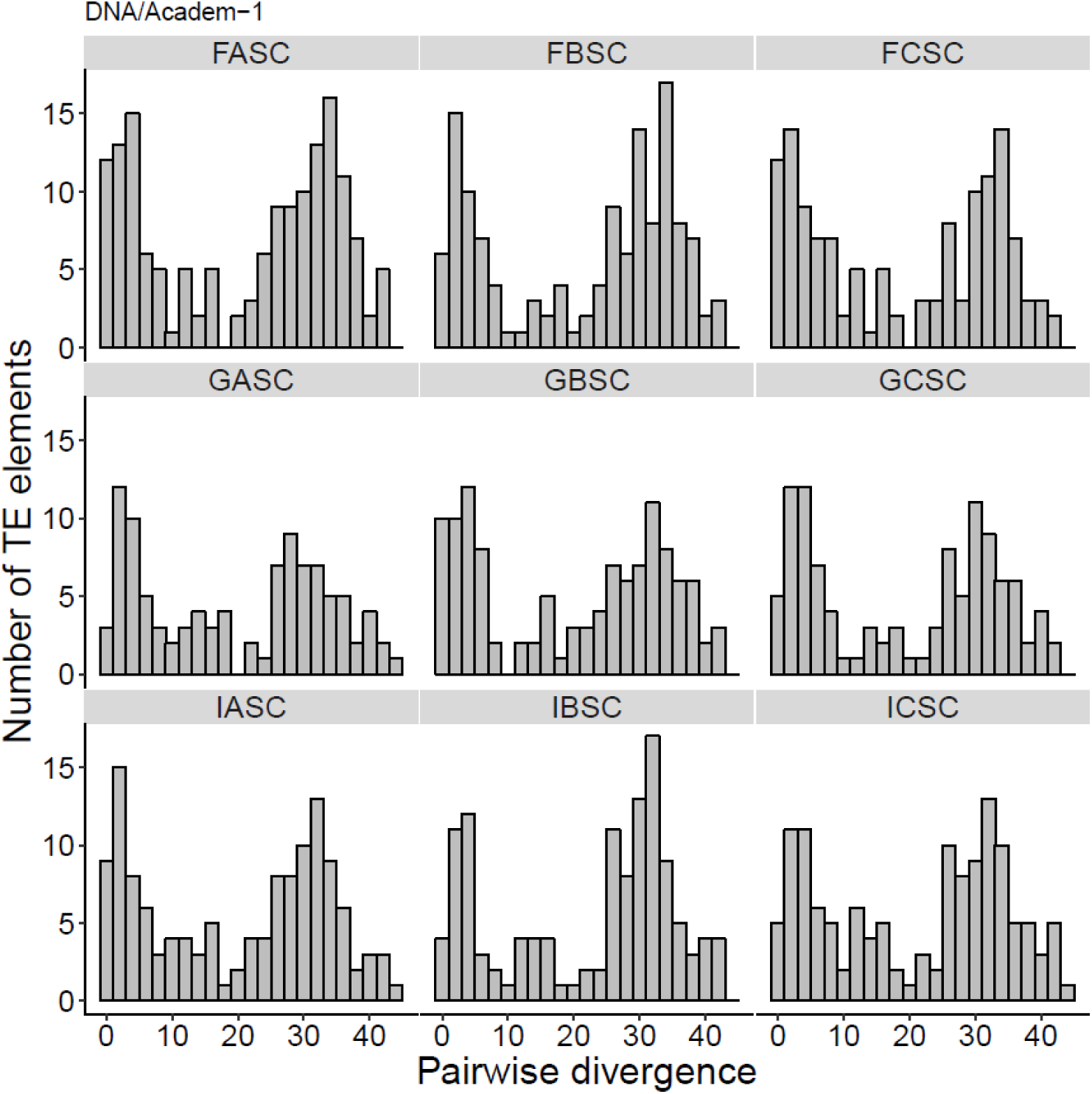
Pairwise divergence of DNA/Academ-1 for all nine reference genomes of *D. magna* originally collected from Finland (FASC, FBSC, and FCSC), Germany (GASC, GBSC, and GCSC), and Israel (IASC, IBSC, and ICSC).

**Figure S4D.**
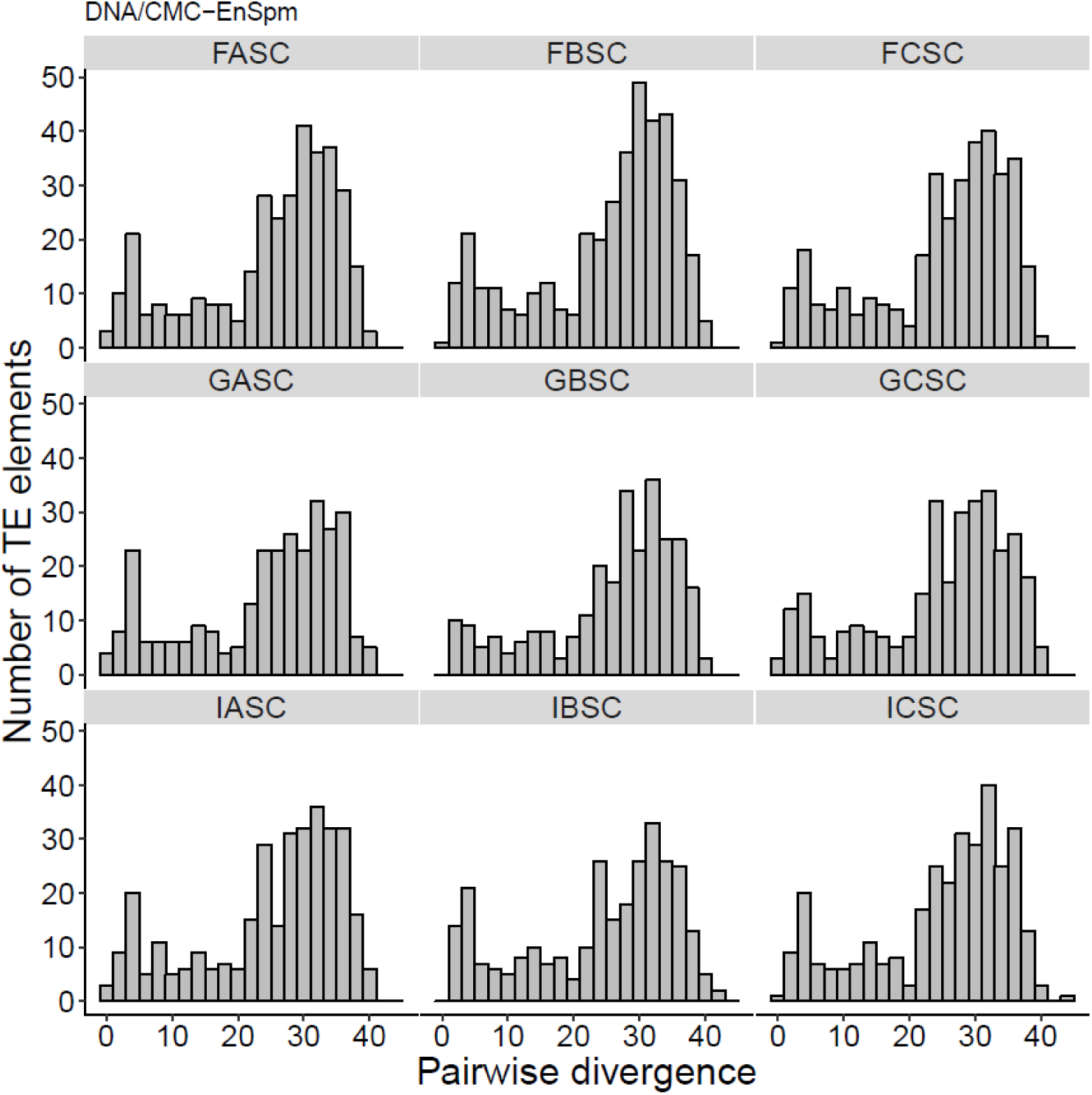
Pairwise divergence of DNA/CMC-EnSpm for all nine reference genomes of *D. magna* originally collected from Finland (FASC, FBSC, and FCSC), Germany (GASC, GBSC, and GCSC), and Israel (IASC, IBSC, and ICSC).

**Figure S4E.**
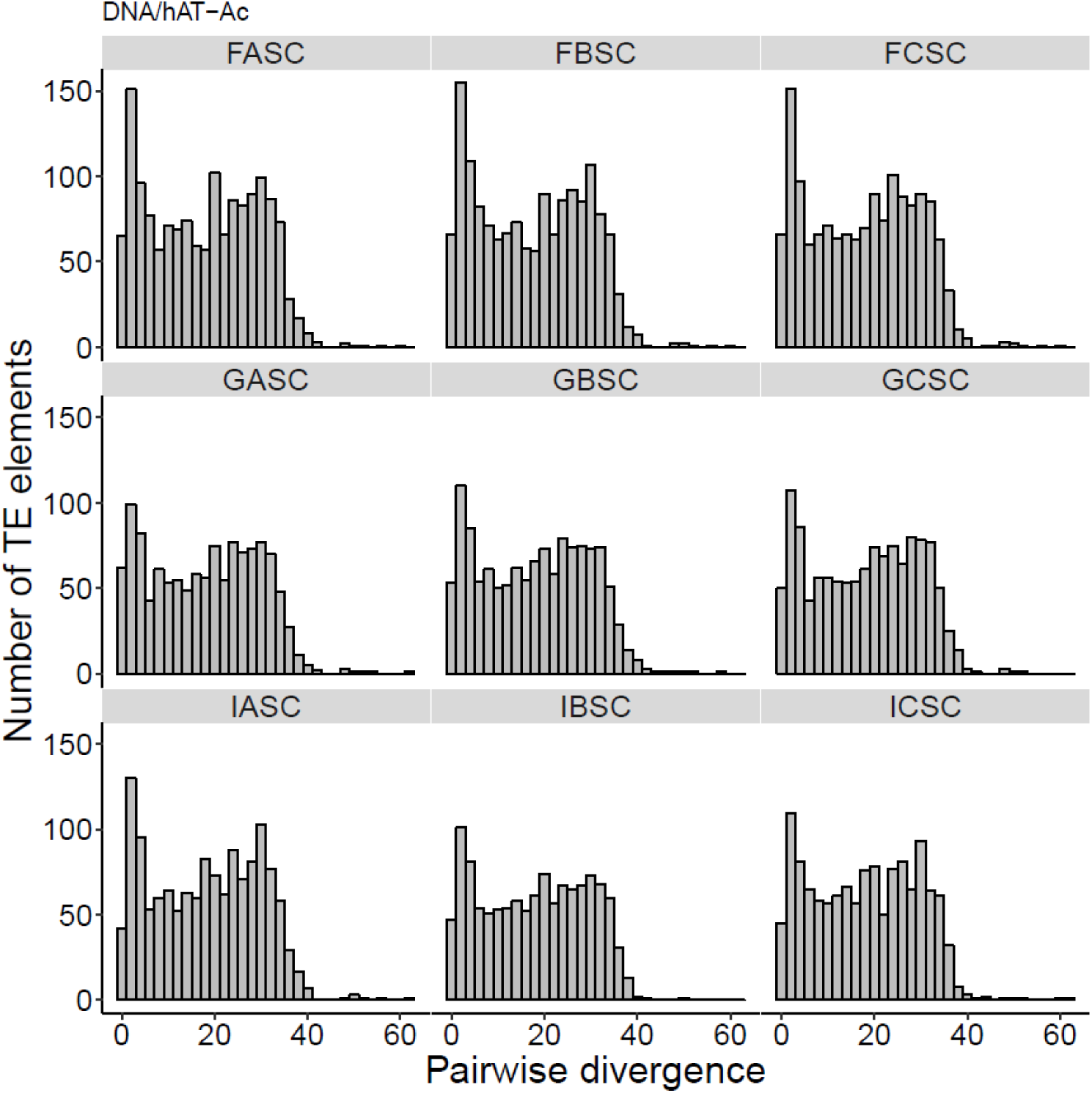
Pairwise divergence of DNA/hAT-Ac for all nine reference genomes of *D. magna* originally collected from Finland (FASC, FBSC, and FCSC), Germany (GASC, GBSC, and GCSC), and Israel (IASC, IBSC, and ICSC).

**Figure S4F.**
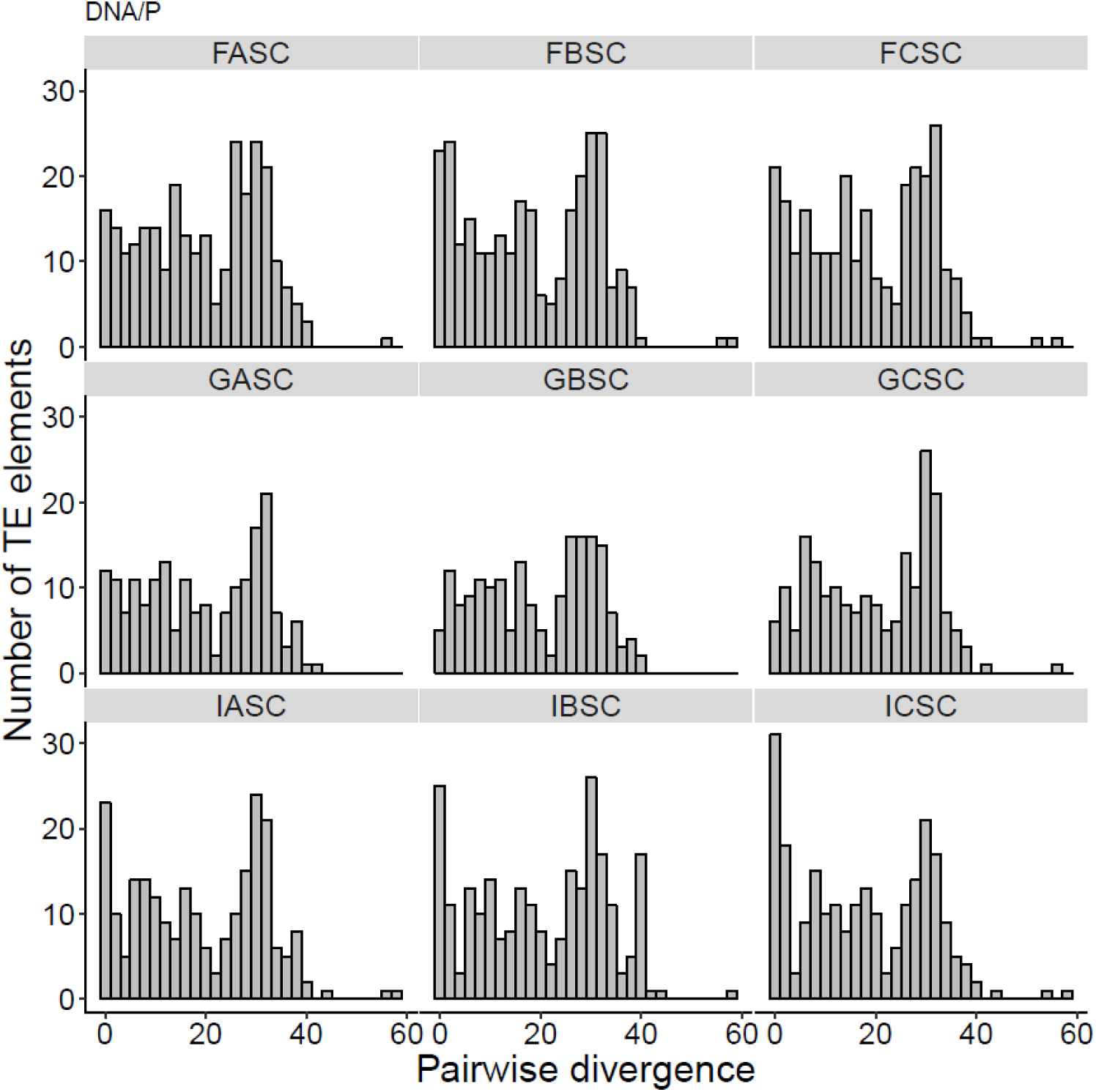
Pairwise divergence of DNA/P for all nine reference genomes of *D. magna* originally collected from Finland (FASC, FBSC, and FCSC), Germany (GASC, GBSC, and GCSC), and Israel (IASC, IBSC, and ICSC).

**Figure S4G.**
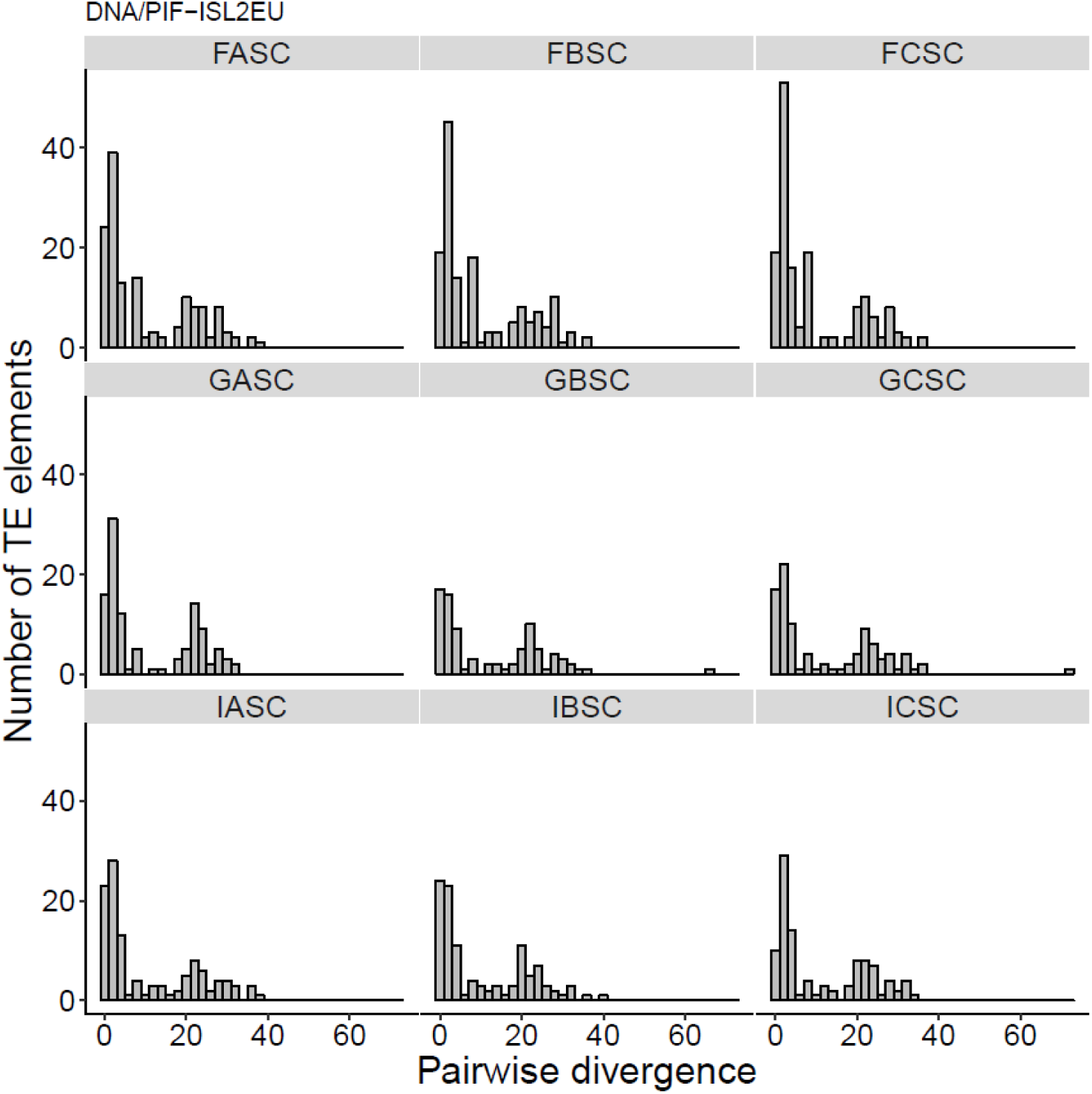
Pairwise divergence of DNA/PIF-ISL2EUfor all nine reference genomes of *D. magna* originally collected from Finland (FASC, FBSC, and FCSC), Germany (GASC, GBSC, and GCSC), and Israel (IASC, IBSC, and ICSC).

**Figure S4H.**
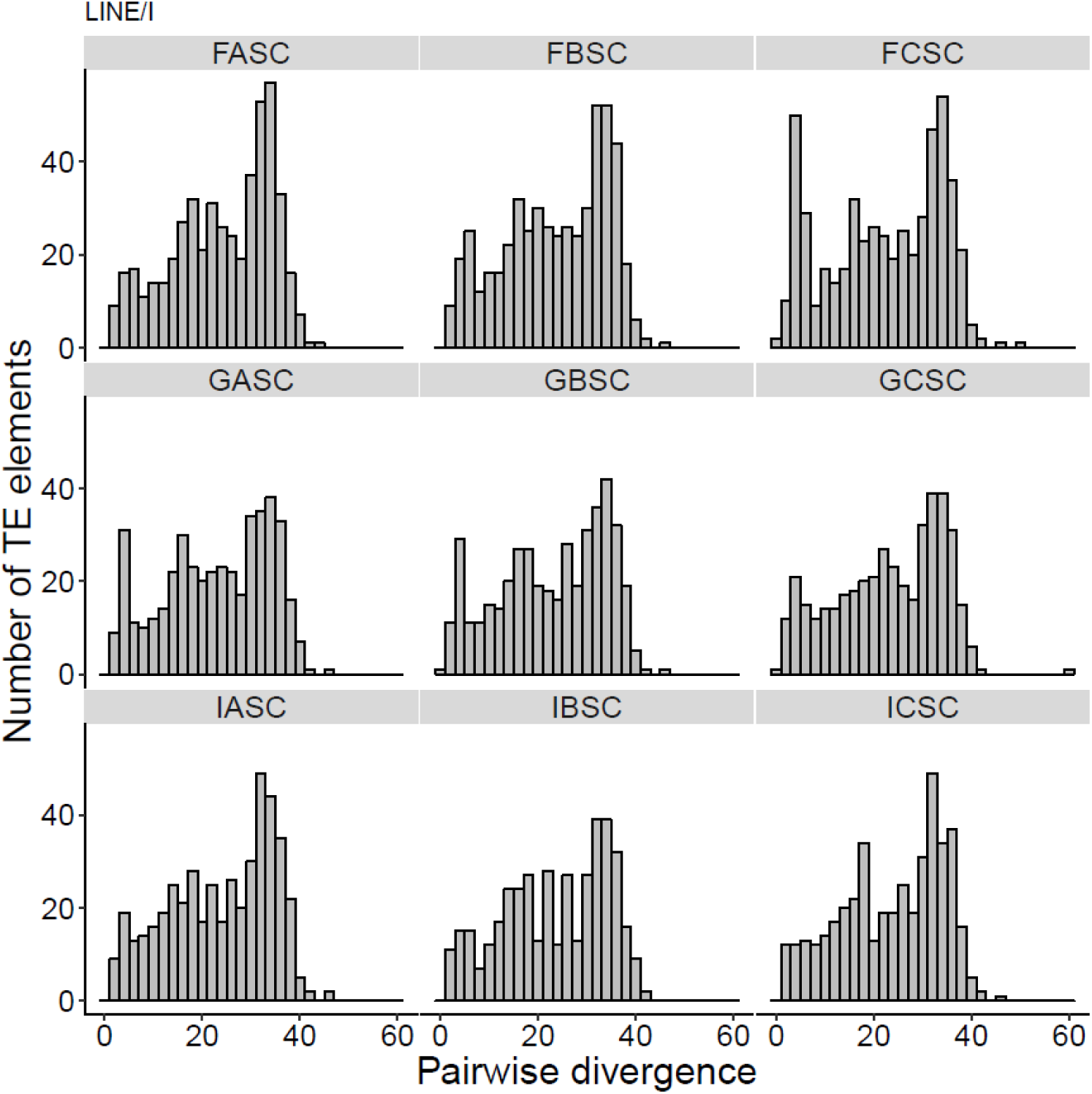
Pairwise divergence of LINE/I for all nine reference genomes of *D. magna* originally collected from Finland (FASC, FBSC, and FCSC), Germany (GASC, GBSC, and GCSC), and Israel (IASC, IBSC, and ICSC).

**Figure S4I.**
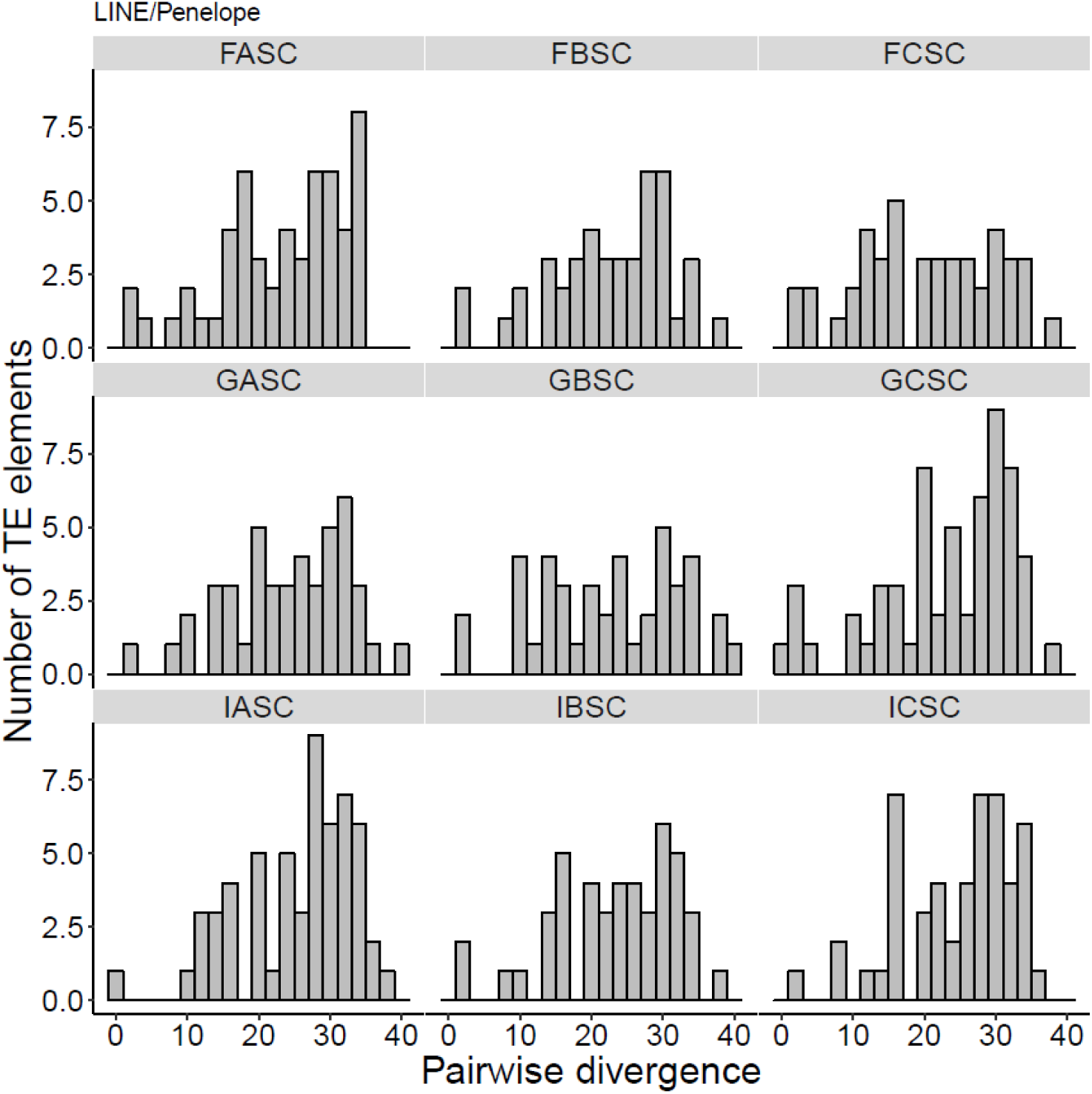
Pairwise divergence of LINE/Penelope for all nine reference genomes of *D. magna* originally collected from Finland (FASC, FBSC, and FCSC), Germany (GASC, GBSC, and GCSC), and Israel (IASC, IBSC, and ICSC).

**Figure S4J.**
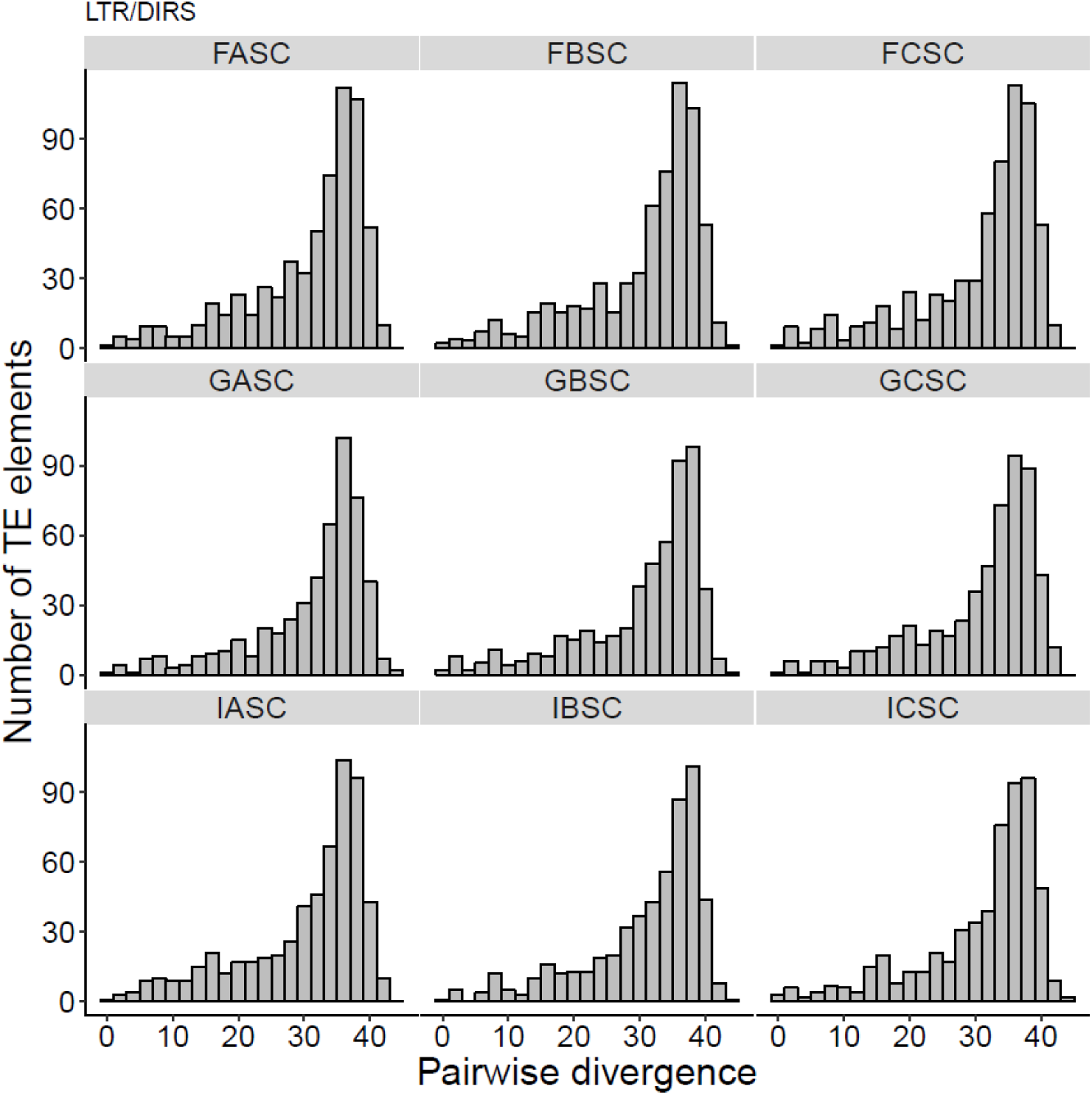
Pairwise divergence of LTR/DIRS for all nine reference genomes of *D. magna* originally collected from Finland (FASC, FBSC, and FCSC), Germany (GASC, GBSC, and GCSC), and Israel (IASC, IBSC, and ICSC).

**Figure S4K.**
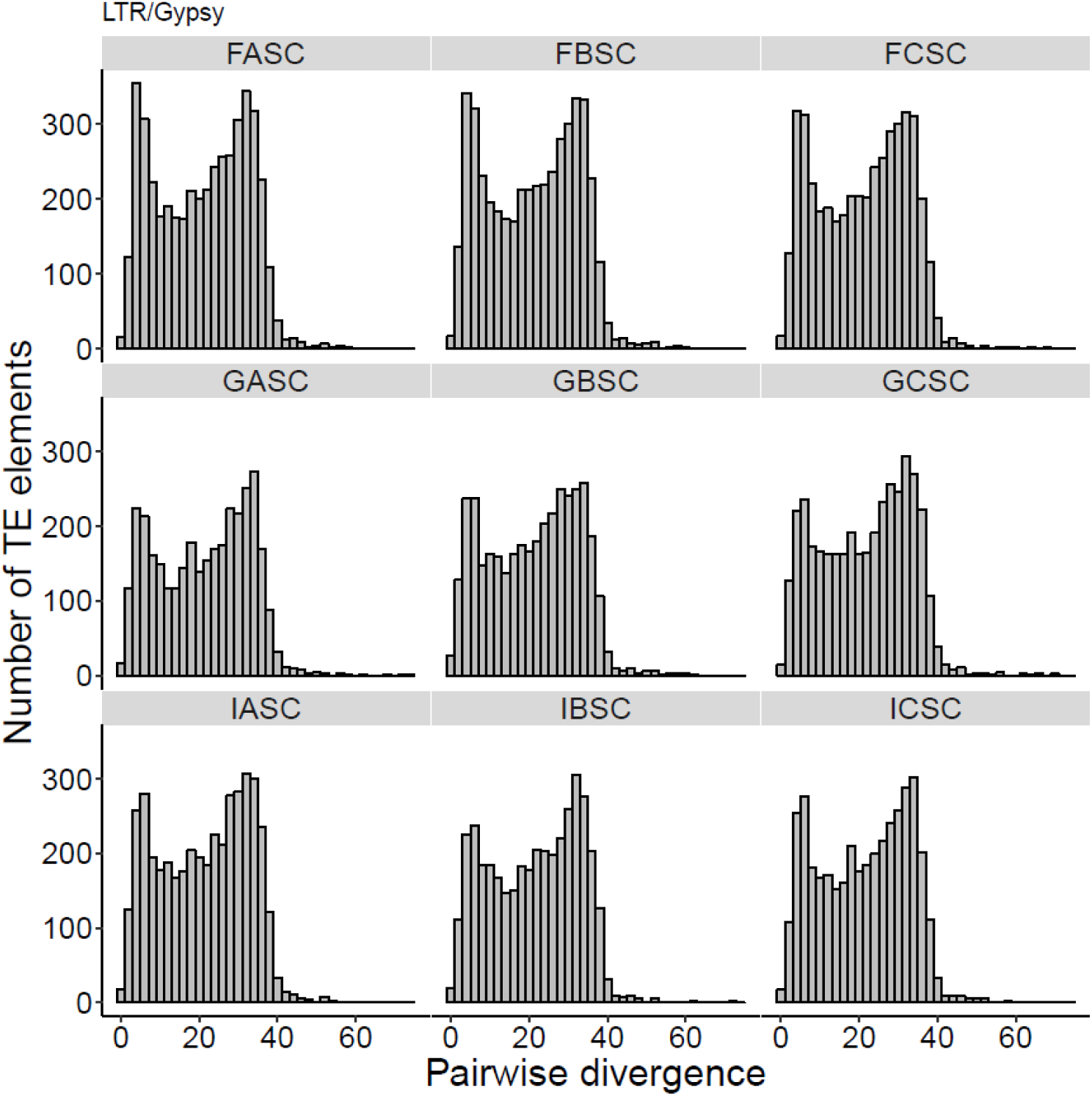
Pairwise divergence of LTR/Gypsy for all nine reference genomes of *D. magna* originally collected from Finland (FASC, FBSC, and FCSC), Germany (GASC, GBSC, and GCSC), and Israel (IASC, IBSC, and ICSC).

**Figure S4L.**
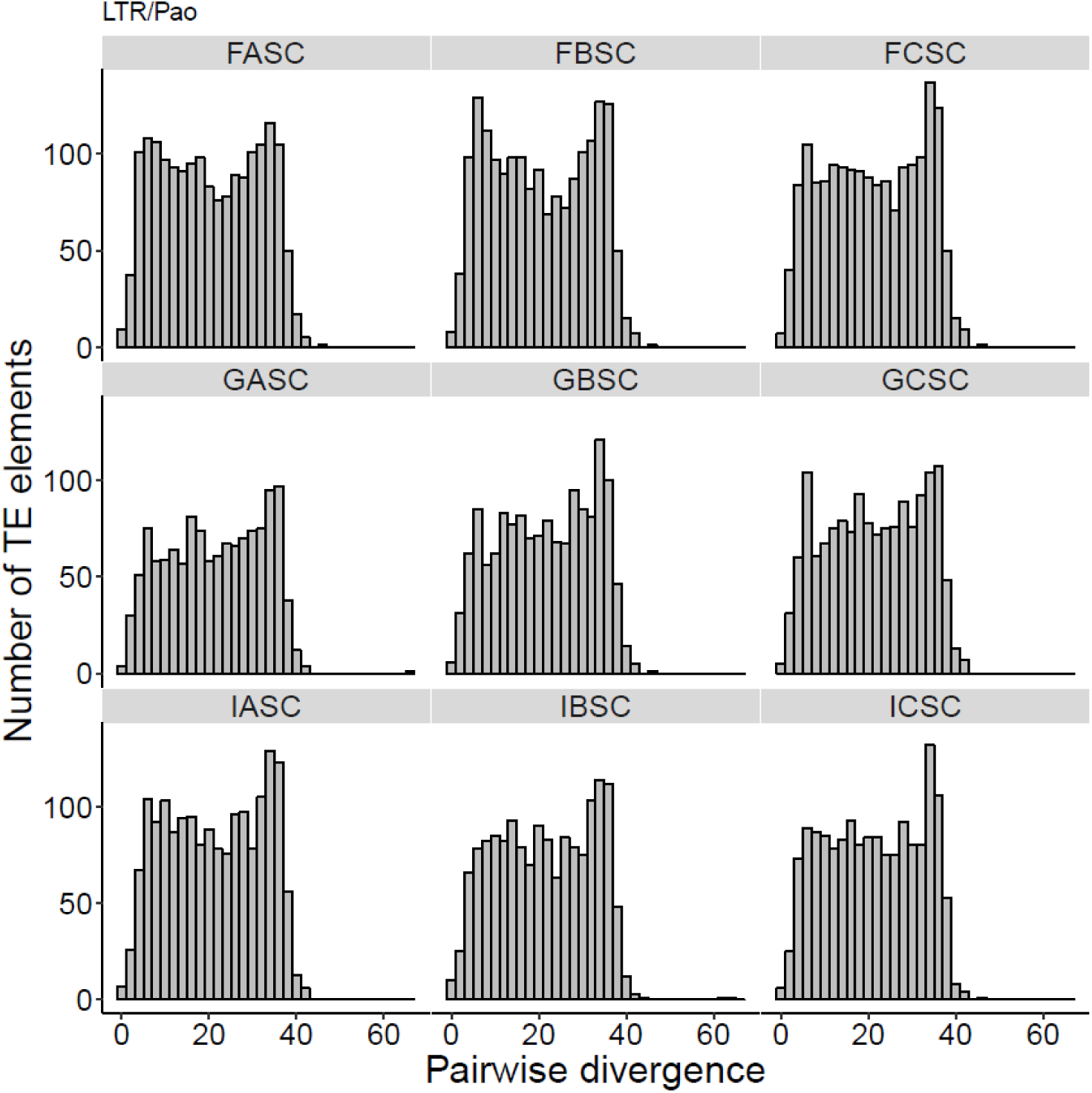
Pairwise divergence of LTR/Pao for all nine reference genomes of *D. magna* originally collected from Finland (FASC, FBSC, and FCSC), Germany (GASC, GBSC, and GCSC), and Israel (IASC, IBSC, and ICSC).

**Figure S5.**
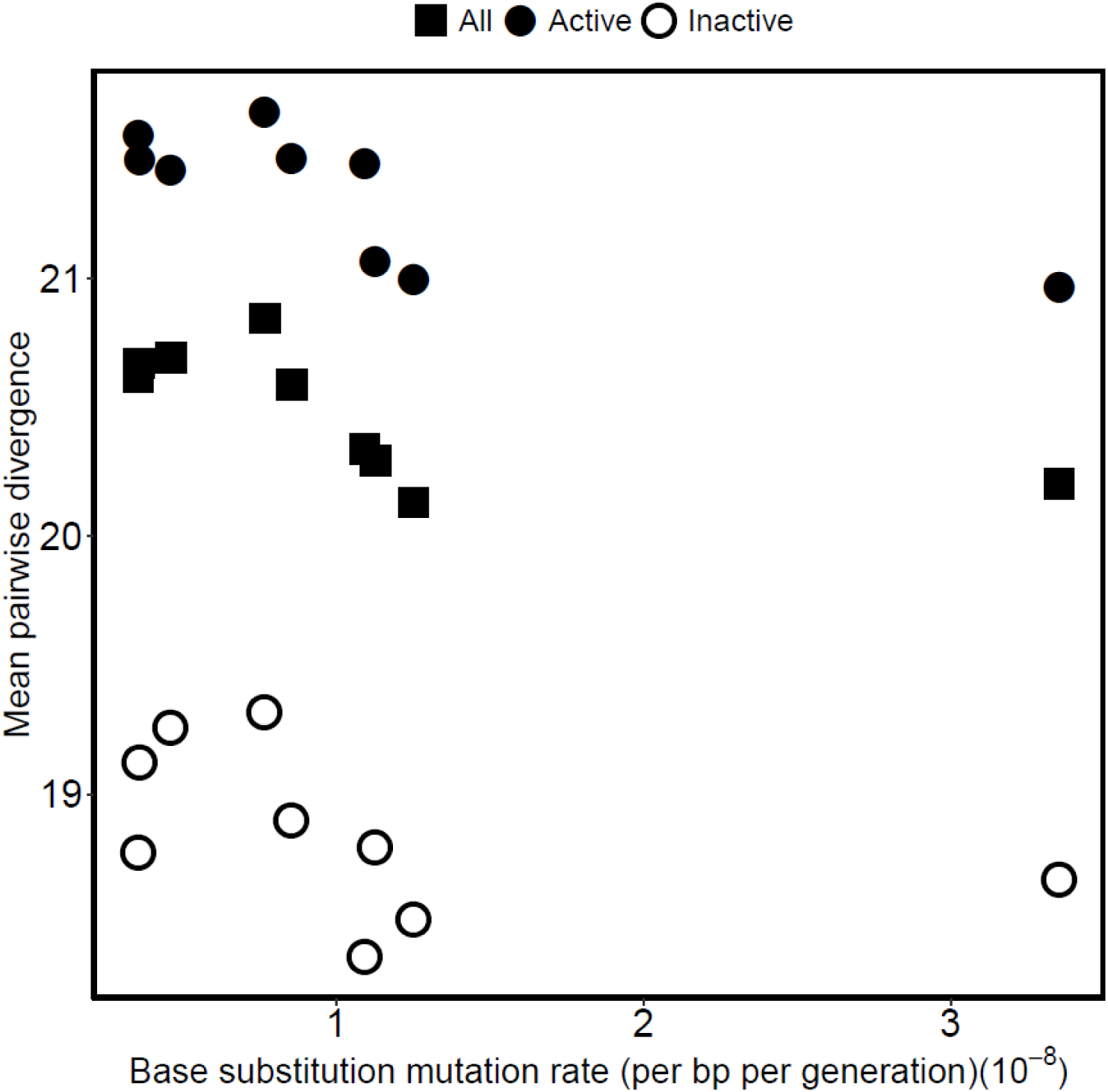
Mean divergence of TE copies (averaged across all TE families) plotted against mean base substitution rate for each starting genotype (filled squares). Mean divergences across only active TE families (based on MA data) are also shown (filled circles) and mean divergence across only inactive TE families (based on MA data) are shown as empty circles.

**Figure S6.**
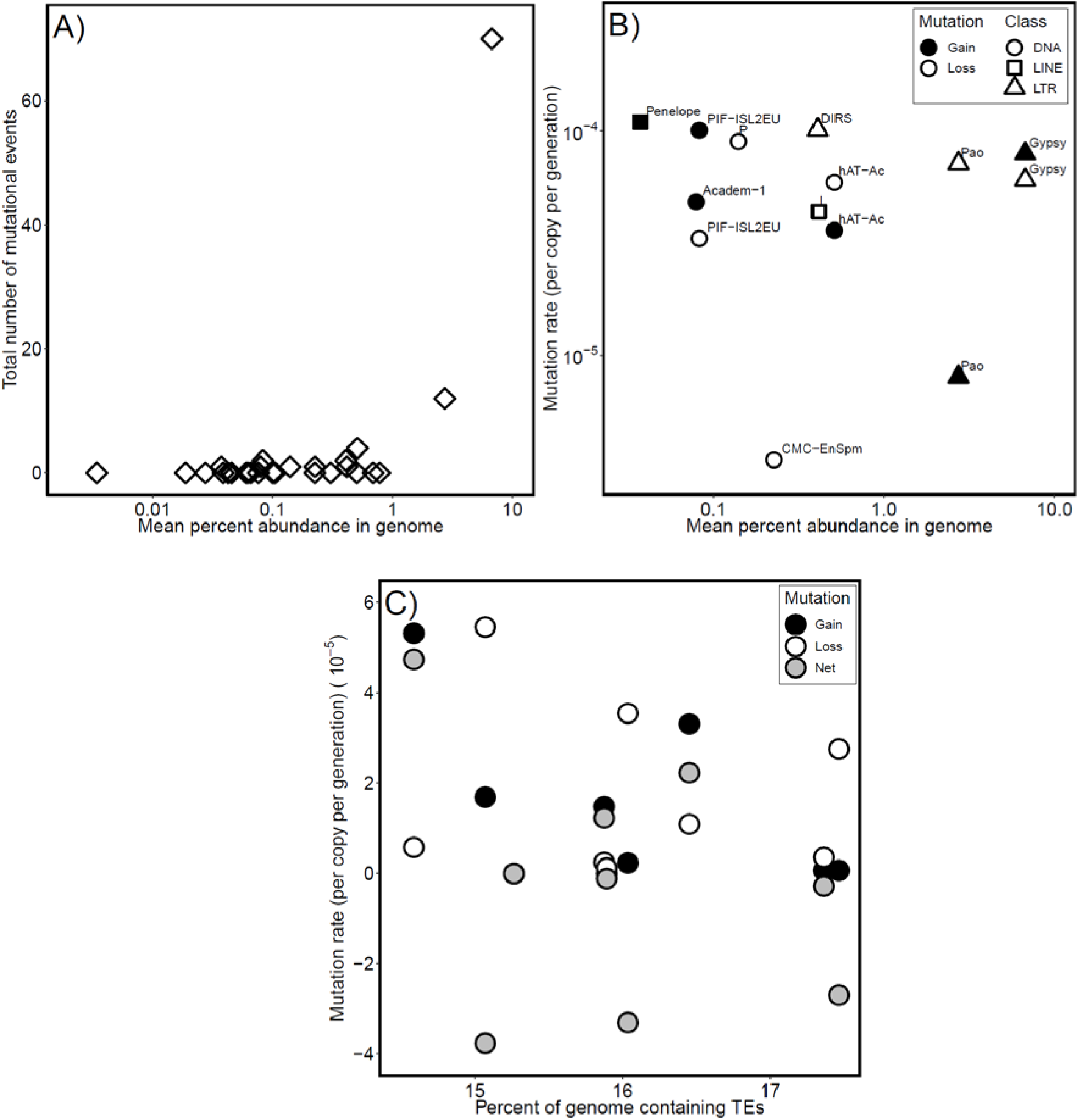
Percent abundance (log scale) of each TE family averaged across genotypes in *D. magna* plotted against (A) number of mutation events for each TE family and (B) gain and loss rates for each active TE family. Gain rates for DNA/CMC-EnSpm, DNA/P, LTR/DIRS, LINE/I and loss rates for DNA/Academ-1, LINE/Penelope are not shown because there were zero mutation events. (C) Percent of the genome occupied by TEs for each assembly plotted against the TE gain (black), loss (white) and net (grey) rates averaged across all families and MA lines. Percent abundance of TEs was estimated using the read mapping approach.

